# Adhesive implant interfaces prevent fibrosis by disrupting mechanobiological feedback

**DOI:** 10.1101/2025.06.01.657311

**Authors:** Mohammad Jafari, Bastien FG Aymon, Yuan Hong, Delaram Shakiba, Guy M Genin, Xuanhe Zhao, Farid Alisafaei

**Affiliations:** Department of Mechanical Engineering, New Jersey Institute of Technology, Newark, NJ; NSF Science and Technology Center for Engineering Mechanobiology; Department of Mechanical Engineering, Massachusetts Institute of Technology, Cambridge, MA; Department of Mechanical Engineering and Materials Science, Washington University in St. Louis, St. Louis; Department of Pathology, Johns Hopkins University School of Medicine, Baltimore, MD; Department of Biomedical Engineering, Washington University in St. Louis, St. Louis, MO; Department of Civil and Environmental Engineering, Massachusetts Institute of Technology, Cambridge, MA

## Abstract

Fibrotic encapsulation around medical implants affects millions of patients annually. Current approaches targeting inflammation or implant material properties have failed clinically, but the mechanical origins of implant-induced fibrosis remain unexplored. Here, we demonstrate that directional imbalance of mechanical forces (“tension anisotropy”) is the primary driver of fibroblast activation at implant-tissue interfaces, and that it can be eliminated through adhesive bonding strategies. Computational modeling reveals a mechanistic basis for successful adhesive anti-fibrotic interfaces: conventional sutured implants generate highly anisotropic stress fields between discrete suture anchor points that activate fibroblasts, while adhesive interfaces distribute forces isotropically, maintaining a mechanical environment that does not activate fibroblasts. *In vivo* experiments from the literature across multiple animal models confirm these predictions: as predicted, adhesive interfaces completely prevent fibrotic capsule formation for up to 12 weeks across diverse organs, while maintaining identical implant composition and geometry compared to sutured controls. Results establish tension anisotropy as a mechanical regulator of implant fibrosis and provide a mechanistic foundation explaining why adhesive interfaces succeed where all previous anti-fibrotic strategies have failed. By addressing the root mechanical cause of fibrosis, this mechanobiology-driven approach may enable a universal approach for preventing fibrosis across all categories of implantable medical devices.

**Significance statement:** Millions of patients suffer from medical device failure due to fibrotic encapsulation, in which a surgically implanted item such as pacemaker leads or a vascular graft loses function by becoming covered with scar tissue. Implants affixed to soft tissues by sutures are especially prone to this form of failure, but implants affixed with a recently invented adhesive are not. We present the discovery that directional imbalance of forces (“tension anisotropy”) drives conversion of healing tissue into scar tissue. Conventional sutured implants create highly anisotropic stress fields between anchor points that activate fibroblasts, while adhesive interfaces distribute forces isotropically, attenuating scarring. This mechanistic insight explains why adhesive implant-tissue interfaces successfully prevent fibrotic capsule formation across multiple animal models and organ systems, where all previous anti-fibrotic approaches have failed. By addressing root mechanical causes of fibrotic remodeling, this discovery provides a pathway for clinical remediation of fibrotic encapsulation.

## Introduction

Fibrotic encapsulation of implanted biomaterials represents one of the most significant challenges in modern medicine, affecting millions of patients annually and compromising the functionality of devices ranging from cardiac pacemakers to continuous glucose monitors.^1^ At the interface between soft tissue and an implanted device, activated fibroblasts assemble a dense, collagen-rich capsule that physically separates the implant from the host tissue, leading to device isolation, reduced efficacy, and often necessitating high-risk revision surgeries. In cardiac devices alone, fibrotic complications require 10,000-15,000 lead extractions annually with significant morbidity and mortality.^2–8^ Despite decades of research targeting inflammation, material properties, and surface modifications, no strategy has successfully prevented fibrotic encapsulation across device types.^9–15^

The cellular mechanisms underlying implant-induced fibrosis center on the fibroblast-to-myofibroblast transition.^16^ In response to biochemical and mechanical cues from the extracellular matrix (ECM),^17–20^ especially ECM stiffness and stress,^21,22^ fibroblasts can transform from quiescent maintainers of ECM homeostasis^23–25^ to highly contractile, synthetic myofibroblasts characterized by α-smooth muscle actin (αSMA) expression and excessive collagen deposition.^26–28^ While this transformation is essential for wound healing, its persistence leads to pathological fibrosis.^29–35^ Current understanding suggests that mechanical factors in the cellular microenvironment play crucial roles in regulating this phenotypic transition, yet the specific mechanical triggers remain incompletely understood.

Recent evidence points to a previously unrecognized mechanical regulator, tension anisotropy: the directionality of mechanical forces, not merely their magnitude, governs fibroblast activation.^36^ Tension anisotropy refers to a condition in which the mechanical tension experienced by a cell is not uniform across directions. In an isotropic stress field, the cell senses equal tension in all directions; in contrast, under anisotropic stress, tension is greater along one axis than another. This directional imbalance is a potent cue that regulates the contractile phenotype of fibroblasts. Although native tissues experience mechanical stresses that typically vary with direction,^37–39^ this is exacerbated at implant-tissue interfaces, particularly around sutured implants where discrete attachment points create highly directional stress fields between sutures. This anisotropic mechanical loading induces long-lasting mechanical memory in fibroblasts through epigenetic modifications, causing persistent activation even after mechanical stimuli are removed.^22^

A potential solution has emerged from our development of adhesive implant-tissue interfaces.^40^ Unlike conventional sutured implants that create mechanical discontinuities, adhesive interfaces form conformal, mechanically continuous bonds with tissue surfaces. In multiple animal models, these adhesive interfaces completely prevented fibrotic capsule formation for up to 12 weeks across diverse organs. Our mechanistic studies now suggest that adhesive bonding eliminates tension anisotropy at the implant-tissue interface, preventing fibroblast activation and fibrotic capsule formation.

The relationship between adhesive interfaces and mechanical anisotropy involves an interplay of cellular and matrix mechanics. During early cell-matrix interactions, fibroblasts extend microtubule-rich protrusions that locally align collagen fibers.^41^ These aligned fibers create anisotropic mechanical environments that, in turn, guide further protrusion growth in a self-reinforcing feedback loop.^36^ At sutured implant interfaces, this process amplifies mechanical anisotropy between discrete attachment points. Conversely, adhesive interfaces may disrupt this feedback by maintaining isotropic stress distributions.

In this study, we demonstrate that tension anisotropy is the primary mechanical driver of implant-induced fibrosis and that its elimination through adhesive bonding prevents fibrotic encapsulation. Using computational mechanics alongside *in vitro* and *in vivo* evidence, we show that adhesive interfaces prevent fibrosis by maintaining isotropic mechanical environments. These findings establish a mechanistic foundation for design strategies that prevent fibrotic encapsulation.

## Results and Discussion

### Tension anisotropy outweighs the effect of ECM stiffness on fibroblast activation

To establish the mechanistic foundation for our central hypothesis that tension anisotropy drives implant-induced fibrosis, we first validated that directional mechanical forces are indeed the primary driver of fibroblast activation. This required demonstrating that tension anisotropy can exert a more powerful influence on fibroblast phenotype than other well-established mechanical factors, including ECM stiffness.

The theoretical basis for this investigation stems from our understanding that fibroblasts regulate tissue mechanics by generating internal contractile forces through their actomyosin machinery.^41–44^ These contractile forces are transmitted to the surrounding ECM, generating mechanical tension within the matrix.^22^ This cell-generated tension not only maintains tissue integrity during homeostasis but also plays a central role in processes such as wound healing, fibrosis, and implant-tissue integration.^45^ Upon activation, fibroblasts upregulate their actomyosin contractility and transition into myofibroblasts, a cell type characterized by enhanced force generation and elevated synthesis of ECM proteins such as collagen.^46^

Mirroring our recent discovery that mechanical stress field anisotropy regulates fibroblast activation more powerfully than stress magnitude alone,^36^ we replicated earlier results with our established micropatterning system that enables independent control of tension magnitude (through ECM stiffness) and tension anisotropy (through cell geometry) at the single-cell level. We studied fibroblasts with equal spreading area cultured on circular and rectangular adhesive islands micropatterned onto deformable polyacrylamide hydrogels of tunable stiffness (9, 18, and 36 kPa). This system enabled us to independently assess how the magnitude and anisotropy of tension regulate fibroblast activation (**Fig. 1**). As fibroblasts contract, they deform the ECM, and the ECM, in turn, resists this contraction. The magnitude of the resulting tension increases with ECM stiffness, as stiffer matrices generate greater resistance to cell-generated forces. The anisotropy of the tension field, however, is governed by cell geometry. Our 3D traction measurements (**Fig. S1**),^36^ together with previous 2D studies,^47,48^ have shown that circular cells experience a planar isotropic stress field, with similar levels of tension in all directions. In contrast, elongated rectangular cells generate an anisotropic stress field, with elevated tension along their long axis compared to the transverse direction.

**Figure 1.**
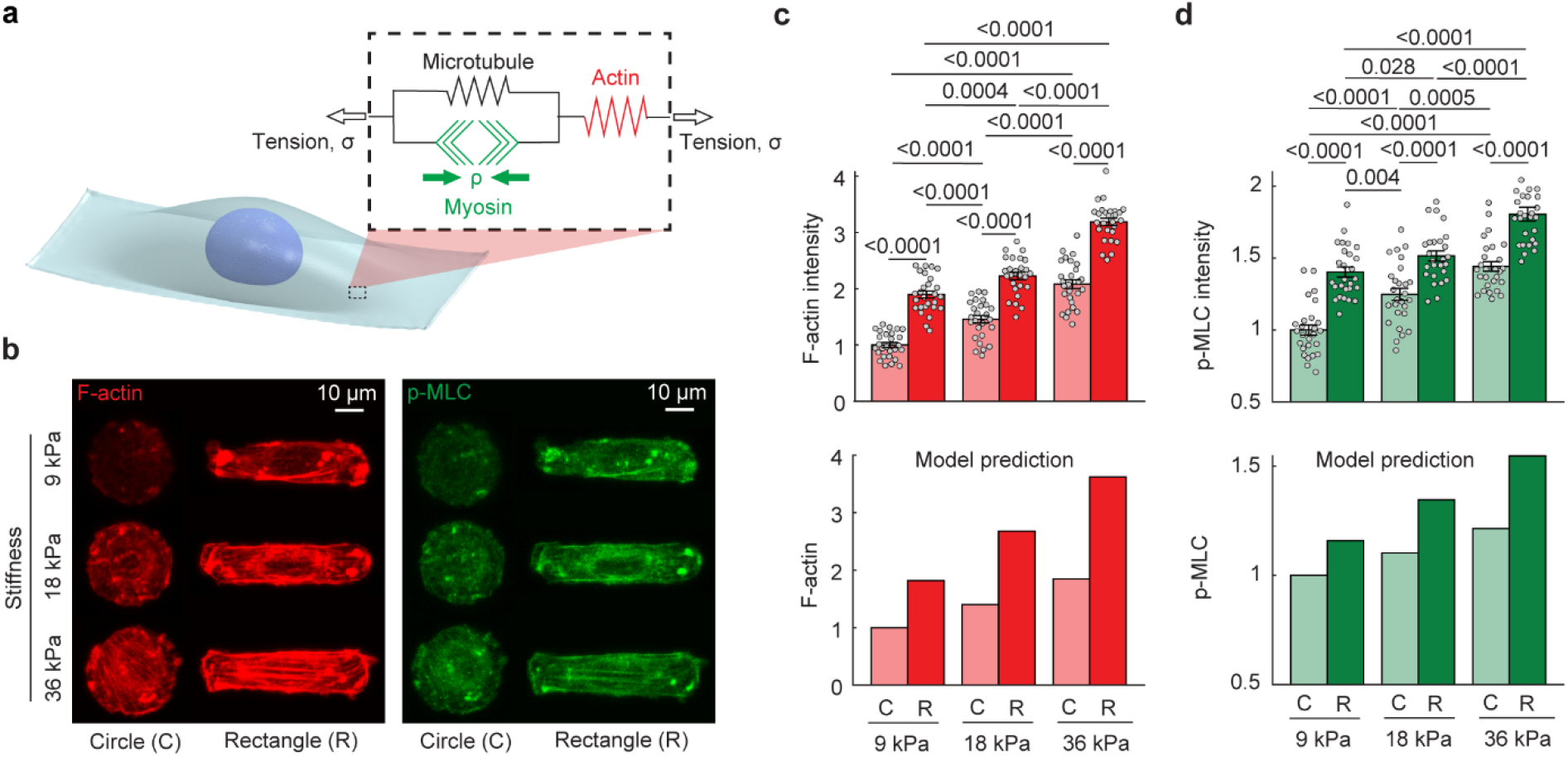
Tension anisotropy enhances fibroblast activation beyond the effects of ECM stiffness at the single-cell level. **a**, Schematic of the theoretical model in which the cytoskeleton is treated as a continuum of representative volume elements (RVEs). Each RVE comprises a contractile myosin element, a passive element mimicking microtubules (arranged in parallel), and another passive element representing actin filaments (arranged in series). Myosin contraction generates an internal force that compresses the microtubule element and stretches the actin element, recapitulating cytoskeletal force transmission. The model is used to predict myosin and F-actin levels in response to changes in cell shape and ECM stiffness. **b**, To test the model prediction, fibroblasts were confined to micropatterned circular (isotropic) or rectangular (anisotropic) patches of identical area on deformable hydrogels with tunable stiffness (9, 18, or 36 kPa), enabling independent control of both ECM stiffness and tension anisotropy to assess their respective contributions to fibroblast activation. **c-d**, As predicted by the model, fibroblasts cultured on rectangular islands exhibited significantly higher levels of F-actin and p-MLC compared to circular cells at each stiffness level, indicating elevated actomyosin contractility driven by tension anisotropy. Notably, fibroblasts on rectangular islands on 9 kPa substrates showed greater activation than circular cells on 18 kPa substrates, demonstrating that directional anisotropy of tension can exert a stronger influence on fibroblast activation than ECM stiffness. In panels **c-d**, *n* = 27 for each group. Panels **c-d** were reproduced from our published data in ref.^*36*^The height of the bars and the error bars indicate the mean and the standard error, respectively. Statistical analysis was performed using two-way ANOVA followed by multiple comparison tests using the Holm-Sidak method.

Our theoretical cell model (implemented into a 3D finite element framework) treats the cytoskeleton as a continuum of representative volume elements (RVEs), each containing three primary elements: myosin, microtubules, and actin filaments (**Fig. 1a**). Central to the model is a feedback mechanism: cells respond to increasing mechanical tension by promoting actomyosin contractility, and this response is amplified by both the magnitude and anisotropy of the tension:^36^

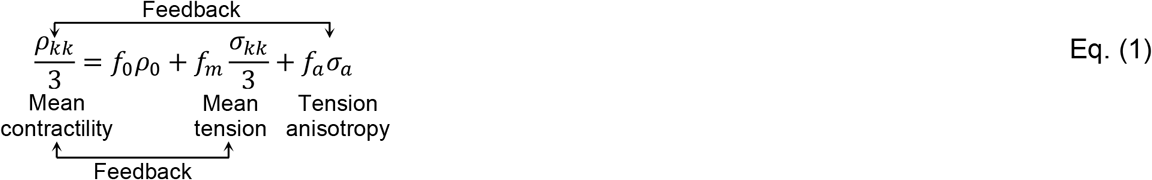

where *ρ*_*kk*_*/* 3 = (*ρ*_11_ + *ρ*_22_ + *ρ*_33_) */* 3 is the average of contractile stress generated by the myosin element in all three directions, indicating overall cell contractility. Similarly, *σ*_*kk*_*/* 3 = (*σ*_11_ + *σ*_22_ + *σ*_33_) */* 3 is the average of tension in all three directions. Here, *σ*_*a*_is tension anisotropy, representing the difference between the maximum and intermediate principal stresses (see SI for more details). Also, *ρ*_0_ denotes the baseline cell contractility in the absence of ECM. Furthermore, *f*_0_, *f*_m_, and *f*_*a*_are model feedback parameters governing the tension-dependent feedback mechanism between cell contractility *ρ* and tension *σσ* (see, SI). As a result of this feedback mechanism, wherein cell actomyosin contractility increases with both the magnitude and directional anisotropy of tension, the model predicted that actomyosin levels rise with increasing ECM stiffness and with increased cell elongation (**Fig. 1c,d**).

Our experimental results, previously published in ref.^36^and reproduced here for context, closely aligned with theoretical predictions and established three key findings. First, fibroblasts increased their actomyosin contractility with ECM stiffness, as indicated by higher levels of filamentous actin (F-actin) and phosphorylated myosin light chain (p-MLC) in both circular and rectangular geometries (**Fig. 1b-d**). This confirmed that tension magnitude alone is sufficient to promote fibroblast activation, consistent with decades of prior research. In the context of implants, this would imply that the mechanics of an implant would directly drive fibroblast activation. However, the observation that suturing and adhesive repair have dramatically different outcomes is at odds with this observation.

Second, at each stiffness level, rectangular cells consistently exhibited higher levels of F-actin and p-MLC than circular cells (**Fig. 1b-d**). This demonstrates that tension anisotropy enhances fibroblast activation independently of ECM stiffness, a finding that directly supports our hypothesis that directional force imbalances drive implant fibrosis. Most critically for our hypothesis, we observed that tension anisotropy can outweigh the effects of ECM stiffness within physiological ranges. Fibroblasts on rectangular islands on soft 9 kPa ECMs exhibited significantly higher activation than circular cells on 18 kPa ECM (**Fig. 1c,d**). This demonstrates that, within the physiological stiffness range,^49,50^anisotropy of tension can have a more pronounced effect on fibroblast activation than ECM stiffness.^36^

These single-cell results provide the mechanistic foundation for a hypothesis that eliminating tension anisotropy at implant-tissue interfaces can prevent fibroblast activation and subsequent fibrosis. If directional force imbalances are indeed more potent activators than matrix stiffness, a factor that has dominated biomaterial design efforts, then strategies targeting tension anisotropy represent a paradigm shift in anti-fibrotic implant development. The next critical test of this hypothesis required demonstrating that these principles hold at the tissue scale, where multiple cells interact within complex mechanical environments that more closely approximate implant-tissue interfaces.^36^

### Tension anisotropy at the tissue level governs fibroblast activation

Having established that tension anisotropy regulates fibroblast activation at the single-cell level (**Fig. 1**), we next investigated whether this principle holds at the tissue scale. To do this, we studied engineered cruciform-shaped tissues. These tissues consist of four arms, with each arm anchored at its end (**Fig. S2**). As fibroblasts within the tissue contract, they generate spatially heterogeneous stress fields arising solely from the geometry of the construct. Cells located near the center of the tissue and close to the arm ends experience equibiaxial, isotropic tension (equal in all in-plane directions), while cells in the midsection of the arms experience anisotropic, uniaxial tension (**Fig. 2a, S2**). This platform enabled us to examine how spatial variations in tension anisotropy (within the same tissue and under uniform culture conditions) impact local patterns of fibroblast activation.

**Figure 2.**
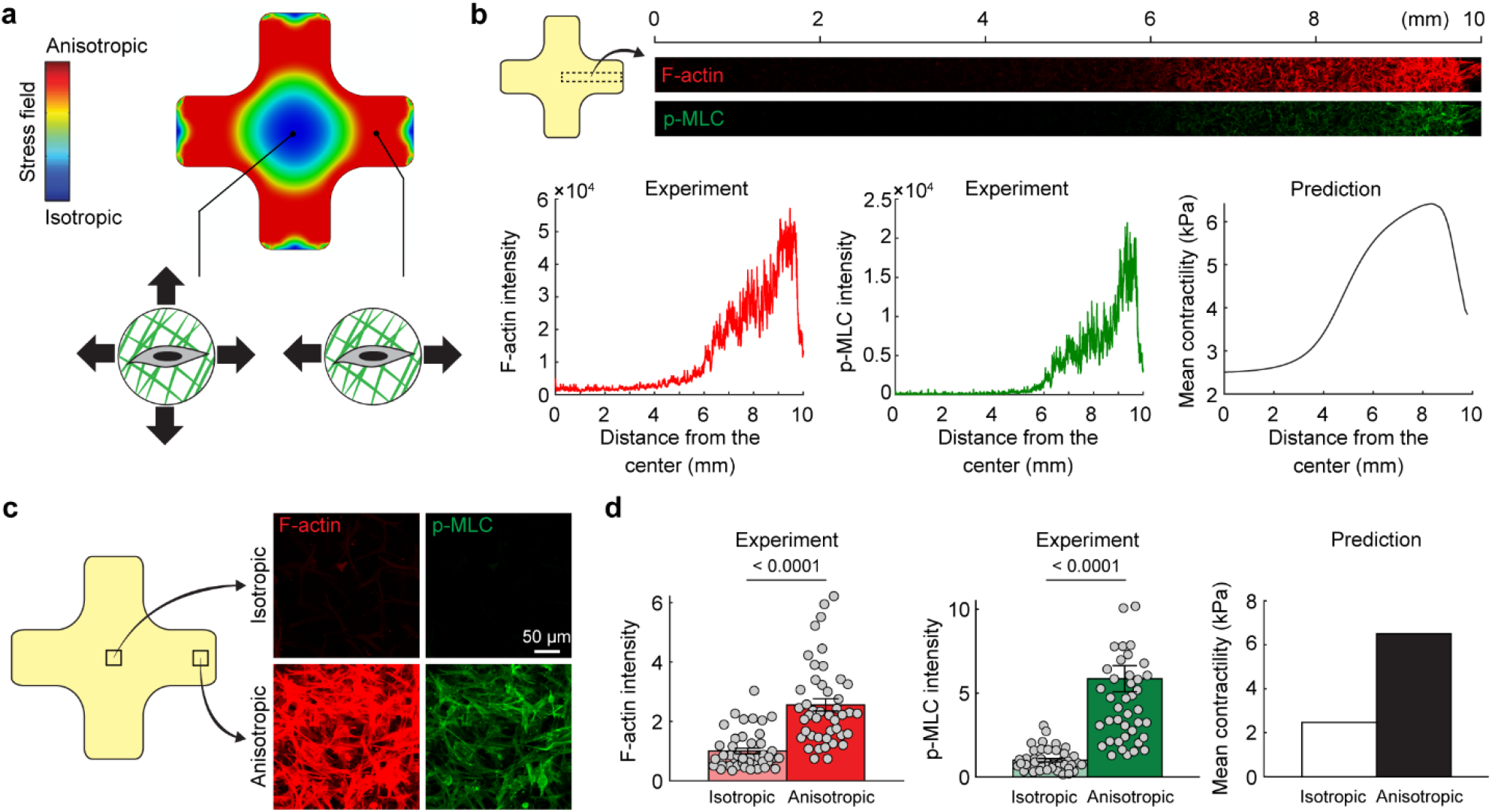
Tension anisotropy, rather than stress magnitude, governs fibroblast activation at the tissue level. **a**, Cruciform-shaped fibroblast-collagen tissues were anchored at the ends of each arm. As fibroblasts contracted, spatially heterogeneous stress fields emerged, with cells in the central region and near the arm ends experiencing isotropic tension, while cells in the mid-arm regions were subjected to anisotropic tension. The heat map illustrates the difference between the maximum and intermediate principal stresses within the tissue, representing the level of tension anisotropy *σ*_a_ in the tissue. Low *σ*_a_ values indicate isotropic stress, whereas high *σ*_a_ values reflect strong tensile anisotropy, i.e., tension predominantly oriented in one direction compared to others. **b**, Quantitative imaging of F-actin and p-MLC along a linear axis extending from the center toward the arm ends revealed that fibroblast activation was suppressed in regions of isotropic tension and enhanced in regions of anisotropic tension. **c**, Elevated levels of F-actin and p-MLC in the mid-arm regions confirmed that tension anisotropy promotes fibroblast activation. **d**, Quantification of F-actin and p-MLC per cell in isotropic versus anisotropic regions validated the predictions of the theoretical model, highlighting tension anisotropy as a dominant mechanical regulator of fibroblast activation in 3D tissue constructs. In panels **d**, *n* = 44 for each group. The height of the bars and the error bars indicate the mean and the standard error, respectively. Panel **d** was reproduced from our published data in ref.^*36*^Statistical analysis was performed using the two-sided unpaired Student’s t-test.

We first used our theoretical model to predict how fibroblast activation varies spatially within cruciform tissues. The model predicted that activation levels would be low in the isotropic center, increase progressively toward the mid-arm regions (where cells experience maximal tension anisotropy), and then decrease again near the anchored arm ends, where tension becomes isotropic once more (**Fig. 2b,d**). Interestingly, when we eliminated the anisotropy term from the model (setting the anisotropy feedback parameter *f*_*a*_= 0 in Eq. (1)), the model predicted the opposite trend, with maximal activation in the isotropic center (**Fig. S3**). This reversed prediction is consistent with prestressed or prestrained cell models and classical thermoelastic-based frameworks, which do not account for the role of tension anisotropy.^36^

Our experimental results validated the predictions of the model. We assessed spatial patterns of fibroblast activation by quantifying the level of fibroblast activation markers (F-actin, p-MLC, and αSMA) across the cruciform tissues. Fibroblasts located in the anisotropic mid-arm regions exhibited significantly elevated levels of all activation markers compared to cells in the isotropic regions located at the center and near the arm ends (**Fig. 2c,d, S4**). These findings support and extend our single-cell results, demonstrating that tension anisotropy is a dominant regulator of fibroblast activation not only in isolated cells but also in collective, 3D tissue environments.

This spatially patterned activation was observed across tissues formed in collagen matrices of varying concentrations (**Fig. S5**), and it persisted even in the presence of a matrix metalloproteinase (MMP) inhibitor (**Fig. S6**), indicating that the observed activation patterns are robust and not dependent on ECM remodeling or degradation.

### Adhesive implants reduce stress anisotropy and fibroblast activation at the implant-tissue interface (model prediction)

Having established that anisotropy of the stress field regulates fibroblast activation at both the single-cell (**Fig. 1**) and tissue level (**Fig. 2**), we next tested the central prediction of our hypothesis: that reducing tension anisotropy at the interface between tissue and an implant could reduce fibroblast activation level and, in turn, limit fibrotic capsule formation. This computational analysis directly addresses a mechanistic hypothesis underlying potential improved anti-fibrotic implant design strategy and provides a theoretical foundation for addressing what remains a persistent challenge in long-term implant integration.^51–58^

Our modeling approach was designed to test the fundamental premise that different implant attachment strategies create distinct mechanical environments that either promote or prevent fibroblast activation. Specifically, we hypothesized that conventional sutured attachments generate highly anisotropic stress fields that drive persistent fibroblast activation, while adhesive interfaces distribute mechanical loads more uniformly, creating isotropic stress environments that suppress activation.

#### Computational prediction of stress fields at sutured interfaces

We first used our theoretical model to predict the mechanical stress distribution at the implant-tissue interface under conventional surgical attachment. We simulated a common clinical scenario in which the implant is attached to tissue using discrete sutures at its edges (**Fig. 3a**), a widely used technique for securing soft implants such as surgical meshes and patches. In our finite element simulations, this condition was modeled by fixing the top-left and top-right points of the tissue to the implant surface (**Fig. 3b**).

**Figure 3.**
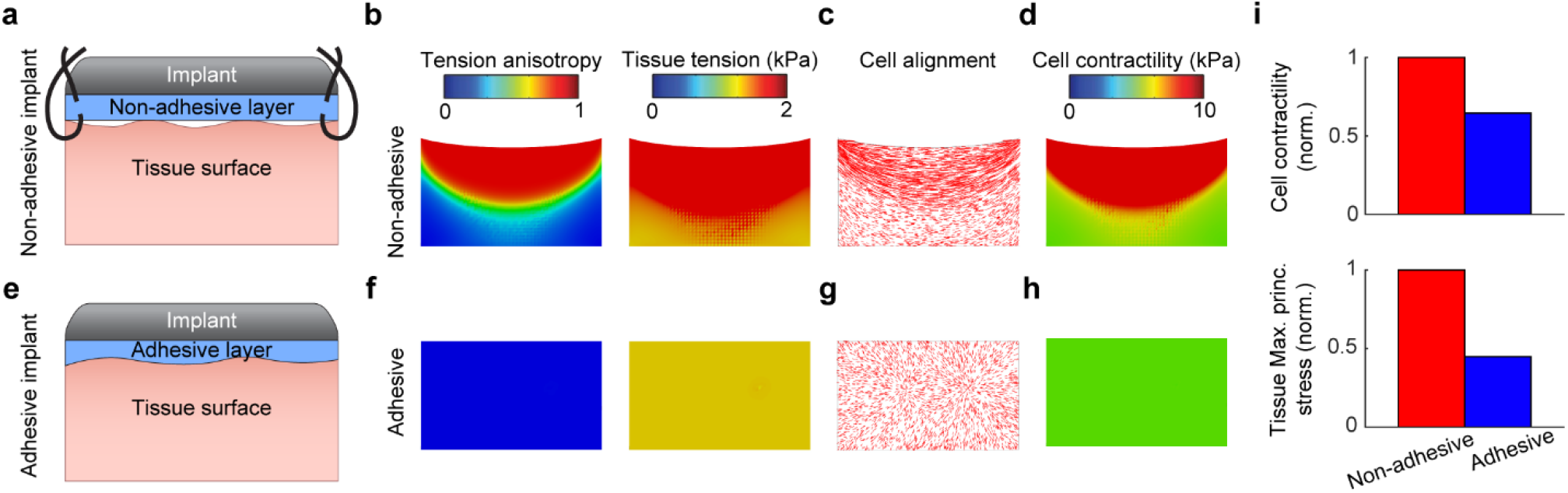
Model prediction for reduced tension anisotropy and fibroblast activation at adhesive implant-tissue interfaces. **a**, Schematic of implant attachment using discrete sutures. **b**, Finite element simulations predicted that sutured implants generated high anisotropic tension *σσ*_aa_ between adjacent sutures, resulting in **c**, aligned fibroblasts along the principal stress direction and **d**, elevated fibroblast activation. **e**, Schematic of an implant with an adhesive interface conformally bonded across the tissue surface. **f**, Simulations showed that adhesive interfaces led to reduced tension anisotropy, **g**, isotropic fibroblast orientation, and **h**, decreased activation levels. **i**, Quantification of fibroblast contractility and tissue tension across the implant–tissue interface reveals significantly lower activation in adhesive compared to sutured conditions.

As fibroblasts contract, the resulting forces generate tension within the surrounding tissue. Due to the free boundary condition in the vertical direction, the upper surface of the tissue, except at the two anchored suture points, is able to contract vertically with minimal resistance. This leads to a surface that exhibits slight inward curvature in the vertical direction, as depicted in our simulations (exaggerated for clarity). In contrast, significant tensile stress develops along the horizontal direction between the sutures, where the tissue is restricted from freely contracting.

#### Critical prediction: localized anisotropic stress drives fibroblast activation

Our model predicted that fibroblasts located between adjacent sutures are exposed to a highly anisotropic stress field (**Fig. 3b**), with mechanical tension concentrated primarily along the axis connecting the sutures. This directional imbalance in mechanical stress led to elevated fibroblast activation and increased contractility, as predicted by the model (**Fig. 3d,i**). The model also predicted that cells would align along the direction of maximal principal stress (**Fig. 3c**), reinforcing the anisotropic mechanical environment through a positive feedback mechanism consistent with our tissue-level observations.

Collectively, these computational results predicted a localized zone of highly activated fibroblasts near the implant-tissue interface, potentially causing fibrotic encapsulation; exactly the clinical problem our study aims to solve through mechanical intervention rather than biochemical approaches that have consistently failed in clinical translation.

#### Adhesive interfaces eliminate stress anisotropy and suppress predicted activation

In contrast, when we simulated an adhesive interface, where the implant is conformally bonded to the underlying tissue across the entire surface (**Fig. 3e**), the model predicted a significant reduction in stress anisotropy at the interface (**Fig. 3f**). This uniform mechanical coupling fundamentally alters the stress distribution compared to discrete suture attachment.

As a result of this mechanical redistribution, fibroblast activation levels were markedly lower (**Fig. 3h, 3i**), and the cells were no longer primarily aligned along a single direction (**Fig. 3g**), indicating a more isotropic mechanical environment. The model predicted that adhesive implants promote a more uniform stress field at the interface and reduce the extent of fibrosis; a prediction that directly supports the central hypothesis of this study.

#### Mechanistic basis for universal anti-fibrotic strategy

These computational predictions provide the mechanistic foundation for our proposed paradigm shift in implant design. Rather than targeting downstream inflammatory or biochemical processes, approaches that have consistently failed in clinical translation, our modeling suggests that addressing the root mechanical cause of fibrosis through tension anisotropy control could provide a universal solution. The model predicts that this mechanical approach should be effective across diverse implant types and anatomical locations, as it targets the fundamental mechanobiological pathway that initiates fibrotic responses.

In summary, our theoretical predictions suggest that adhesive bonding between implant and tissue can mitigate fibroblast activation by redistributing mechanical stresses more isotropically, thereby limiting the development of fibrotic capsules. These computational results establish the mechanistic rationale for the experimental validation studies that follow, where we test whether eliminating tension anisotropy at real implant-tissue interfaces can indeed prevent fibrosis *in vivo*.

### Reducing tension anisotropy at the implant-tissue interface *in vivo*

We next sought to provide evidence that tension anisotropy at the implant-tissue interface may regulate fibroblast activation and contribute to driving fibrotic capsule formation *in vivo*. We have recently developed a bioadhesive hydrogel that enables robust, long-term attachment to diverse organs and tissues.^40^This adhesive forms a conformal, mechanically continuous interface, in contrast to conventional methods where implants are secured using discrete sutures at their edges, leaving the majority of the interface mechanically unconstrained.

To compare these two attachment strategies, we used our previously established rodent model of abdominal wall implantation.^40^Implants were introduced either with a non-adhesive interface (**Fig. 4a**), where their adhesive properties were deactivated and they were then fixed to the tissue using sutures, or with an adhesive interface (**Fig. 4b**), where they were covalently bonded to the tissue using the adhesive hydrogel. This model allowed us to evaluate how mechanical attachment, and particularly differences in stress anisotropy, influence fibrosis *in vivo*.

**Figure 4.**
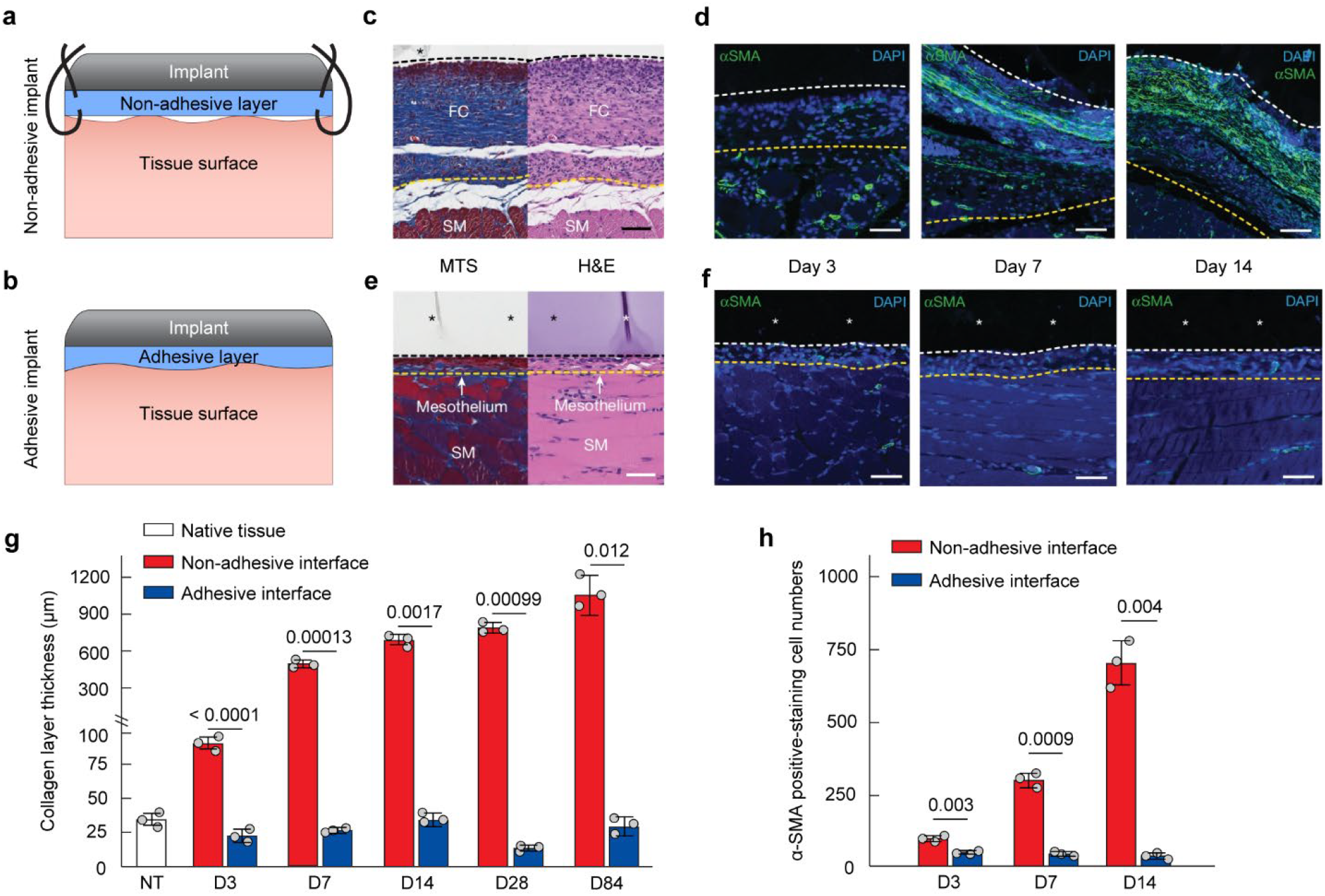
Adhesive implants reduce tension anisotropy and suppress fibrotic encapsulation at the tissue-implant interface. **a,b.** Schematic representation of implant attachment strategies, where (**a**) non-adhesive implants are fixed to tissue using discrete sutures, and (**b**) adhesive implants are bonded conformally to tissue using a bioadhesive hydrogel. **c.** Masson’s trichrome staining (MTS) and Hematoxylin and Eosin Staining (H&E) for tissue collected on Day 14 after implantation on the rat’s abdominal wall showed dense fibrotic capsule formation at the tissue-implant interface of non-adhesive implants. Scale bar 100 μm. **d.** Immunostaining showed elevated α-SMA expression and fibroblast alignment along the implant interface in the non-adhesive condition, with activation levels increasing over time. Scale bars 20 μm for Day 3 and 40 μm for Days 7 and 14. **e.** Adhesive implants exhibited markedly reduced fibrotic encapsulation. Scale bar 50 μm. **f.** α-SMA staining showed reduced fibroblast activation and lack of directional alignment in the adhesive group. Scale bar 20 μm. **g.** Collagen layer thickness at the implant-tissue interface was measured at different time points after implantation. **h.** Quantification of cell numbers in the collagenous layer at the tissue-implant interface over a representative width of 500 µm from the immunofluorescence images on Days 3, 7, and 14 after implantation. In panels **c-f**, asterisks indicate the implant, black or white dashed lines indicate the implant-tissue interface, and yellow dashed lines indicate the mesothelium-fibrous capsule (for non-adhesive implant) or the mesothelium–skeletal muscle interface (for adhesive implant). SM: skeletal muscle, FC: fibrous capsule. In panels **g,h**, n = 3 for each group. NT: native tissue. The height of the bars and the error bars indicate the mean and the standard error, respectively. Statistical analysis was performed using the two-sided unpaired Student’s t-test. Images in **c-f** were taken from our published study in ref ^*40*^. Data in **g,h** were extracted from our published study in ref^*40*^.

Histological analysis of explanted tissues 14 days post-implantation revealed striking differences between the two groups. In the non-adhesive condition, we observed the formation of a dense fibrotic capsule under the implant (**Fig. 4c,g**), with fibroblasts aligned along the long axis of the interface. Immunostaining for α-smooth muscle actin (α-SMA) revealed high levels of fibroblast activation in the fibrotic region (**Fig. 4d**), with activation increasing progressively from day 3 to day 7 and 14 (**Fig. 4h**). These results are consistent with our model predictions that anisotropic stress fields between sutures drive fibroblasts toward a contractile, myofibroblast phenotype.

In contrast, implants with adhesive interfaces exhibited dramatically reduced fibrotic encapsulation (**Fig. 4e,g**). Fibroblasts in the peri-implant region displayed significantly lower α-SMA expression (**Fig. 4f,h**), and cells were not preferentially aligned along the implant interface, both consistent with model predictions. Notably, these differences emerged despite identical implant geometry and material composition, indicating that the observed reduction in fibrosis was not driven by biochemical differences, but rather by the mechanical distinction between adhesive and non-adhesive interfaces.

Together, these results provide *in vivo* evidence that adhesive implants, which reduce tension anisotropy by distributing mechanical loads more uniformly, can effectively suppress fibroblast activation and minimize fibrosis at the implant-tissue interface.

## Conclusions

The formation of a dense fibrotic capsule around implants remains a major clinical challenge, contributing to high device failure rates. Our findings offer a mechanistic understanding of how mechanical design strategies can enhance implant biocompatibility and long-term integration. Specifically, reducing stress anisotropy at the implant-tissue interface through adhesive bonding promotes mechanical continuity, which in turn mitigates the fibrotic response.

Our multi-scale investigation reveals that tension anisotropy serves as a key regulator of fibroblast activation, operating through a self-reinforcing mechanobiological feedback loop. At the single-cell level, fibroblasts experiencing anisotropic tension fields exhibit enhanced actomyosin contractility and myofibroblast differentiation, even on relatively soft substrates. This finding challenges the prevailing paradigm that ECM stiffness alone governs fibroblast activation, demonstrating instead that the directionality of mechanical forces can override stiffness cues within physiological ranges. The tissue-level experiments using cruciform constructs illustrate this principle, showing spatially patterned activation corresponding to regions of tension anisotropy rather than stress magnitude.

Our analyses of implant-tissue interfaces reveal why conventional sutured attachments inevitably fail to prevent fibrosis: discrete mechanical anchoring points create zones of highly anisotropic stress that perpetually activate resident fibroblasts. In contrast, adhesive interfaces distribute mechanical loads uniformly across the entire contact area, maintaining an isotropic stress environment that does not trigger pathological activation (**Fig. 5**). The dramatic reduction in α-SMA expression and collagen deposition observed at adhesive interfaces *in vivo* validates our mechanistic model and demonstrates the therapeutic potential of this approach.

**Figure 5.**
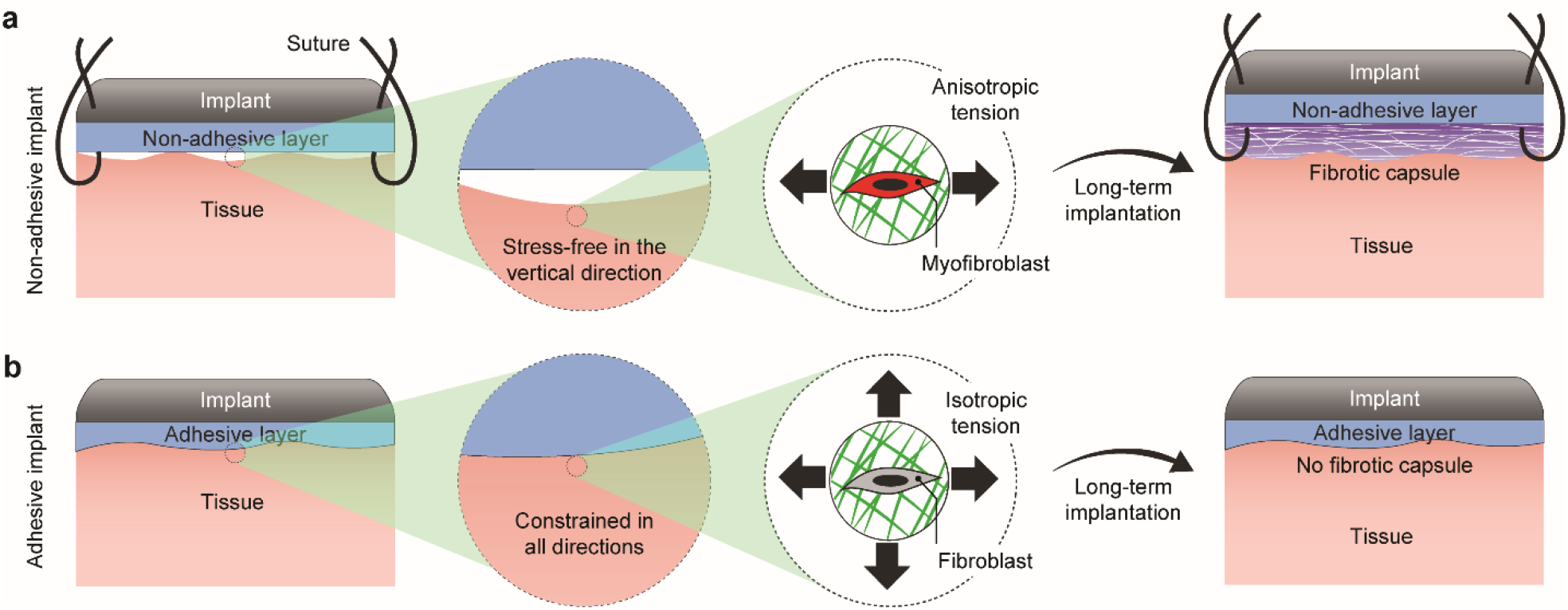
Adhesive implants reduce tension anisotropy and prevent fibrotic encapsulation. **a.** In conventional implants (e.g., sutured), the lack of mechanical attachment at the tissue-implant interface leaves the tissue stress-free in the vertical direction, resulting in anisotropic tension on cells. This can cause long-term activation of fibroblasts into myofibroblasts, ultimately driving excessive collagen deposition and fibrotic capsule formation. **b.** In contrast, adhesive implants form conformal contact with the surrounding tissue, constraining tension uniformly and generating isotropic mechanical fields. This isotropic environment suppresses cellular activation and effectively prevents fibrotic capsule formation during long-term implantation.

Results represent a new approach to anti-fibrotic strategies. Rather than applying biochemical approaches that have consistently failed in clinical translation, our work suggests that addressing the fundamental mechanical drivers of fibrosis may be more effective. Because tension anisotropy appears to be a conserved mechanism across tissue types, adhesive interface strategies may provide a universal solution for preventing fibrotic encapsulation across diverse implant categories, from cardiac devices to neural interfaces. Our findings establish clear mechanical design criteria: minimize tension anisotropy through continuous, compliant, conformal bonding rather than discrete mechanical attachments.

Several limitations merit consideration. While rodent models demonstrate proof-of-concept, translation to humans will require optimization of adhesive formulations for long-term stability of adhesive interfaces. Additionally, the potential for adhesive interfaces to impede device removal must be evaluated.

In conclusion, our study establishes tension anisotropy as a fundamental mechanical regulator of implant-induced fibrosis and demonstrates that adhesive interfaces can prevent fibrotic encapsulation by maintaining isotropic stress distributions. These findings not only explain why previous anti-fibrotic strategies have failed but also provide a clear path forward for developing implantable devices that resist fibrotic encapsulation. By addressing the root mechanical cause of implant fibrosis rather than its downstream consequences, adhesive interface technologies may finally enable long-term integration of implanted devices without the burden of fibrotic isolation, potentially improving outcomes for millions of patients worldwide.

## Acknowledgments

This work was funded in part by the NSF Science and Technology Center for Engineering Biology (CMMI 1548571), the National Institutes of Health (R01AR077793) and the Human Frontier Science Program (HFSP-RGP016/2024).

## Notes

https://github.com/Farid-Alisafaei/Implant-Fibrotic-Capsule

### Competing Interest Statement

The authors have declared no competing interest.

