## Supplementary Information for "Adhesive implant interfaces prevent fibrosis by disrupting mechanobiological feedback"

#### Theoretical cell model

We here present the theoretical cell model, implemented within a finite element framework. The custom code for the finite element simulations is publicly available on GitHub at:

https://github.com/Farid-Alisafaei/Implant-Fibrotic-Capsule

##### Three-dimensional cell model

In the theoretical cell model, the cell is represented as a continuum of representative volume elements (RVEs), each comprising three primary components (Fig. 1a): a contractile myosin element, a compressive microtubule element connected in parallel, and a tensile actin element connected in series. Myosin contraction generates internal forces that compress the microtubule element while simultaneously stretching the actin element, recapitulating the transmission of forces through the cytoskeletal network. Therefore, part of the myosin contractility results in compressive stress in the microtubule, while the rest of that induces tensile stress on the actin and cytoskeleton.

The cell model is formulated based on experimental observations showing that actomyosin contractility increases in response to mechanical tension, whether generated intrinsically by the cell or applied externally.[1–6] This relationship is captured by minimizing $W$, the total free energy of the contractile unit comprising the myosin and the parallel spring element, as described by the following equation:

$$W(\rho,\varepsilon_{\mathrm{c}}) = \frac{1}{3}\,\rho_{kk}\,\varepsilon_{kk}^{(\mathrm{c})} + \tilde{\rho}_{ij}\,\tilde{\varepsilon}_{ij}^{(\mathrm{c})} + \frac{\beta_{\mathrm{v}}}{2}\frac{1}{3}(\rho_{kk}-3\rho_0)^2 + \frac{\beta_{\mathrm{d}}}{2}\tilde{\rho}_{ij}\tilde{\rho}_{ij} - \frac{1}{3}\int_0^{\rho_{kk}}\alpha_{\mathrm{v}}\sigma_{kk}\,d\rho_{kk} - \int_0^{\tilde{\rho}_{ij}}\alpha_{\mathrm{d}}\tilde{\sigma}_{ij}\,d\tilde{\rho}_{ij} + \frac{K_{\mathrm{MT}}}{2}(\varepsilon_{kk}^{(\mathrm{c})})^2 + \mu_{\mathrm{MT}}\tilde{\varepsilon}_{ij}^{(\mathrm{c})}\tilde{\varepsilon}_{ij}^{(\mathrm{c})} - \frac{1}{3}\int_0^{\varepsilon_{kk}^{(\mathrm{c})}}\sigma_{kk}\,d\varepsilon_{kk}^{(\mathrm{c})} - \int_0^{\tilde{\varepsilon}_{ij}^{(\mathrm{c})}}\tilde{\sigma}_{ij}\,d\tilde{\varepsilon}_{ij}^{(\mathrm{c})} \qquad \text{(S1)}$$

where $\rho_{ij}$, $\sigma_{ij}$, and $\varepsilon_{ij}^{(\mathrm{c})}$ are myosin contractility, cytoskeletal stress, and contractile unit strain tensors, respectively (three-dimensional representations of $\rho$, $\sigma$, and $\varepsilon^{(\mathrm{c})}$ in Fig. S7). $\tilde{\rho}_{ij}$, $\tilde{\sigma}_{ij}$, and $\tilde{\varepsilon}_{ij}^{(\mathrm{c})}$ are deviatoric parts of the contractility, stress, and contractile unit strain tensors, and are written as:

$$\tilde{\rho}_{ij} = \rho_{ij} - \frac{1}{3}\rho_{kk}\delta_{ij}$$
$$\tilde{\sigma}_{ij} = \sigma_{ij} - \frac{1}{3}\sigma_{kk}\delta_{ij} \qquad \text{(S2)}$$
$$\tilde{\varepsilon}_{ij}^{(\mathrm{c})} = \varepsilon_{ij}^{(\mathrm{c})} - \frac{1}{3}\varepsilon_{kk}^{(\mathrm{c})}\delta_{ij}$$

The first two terms in equation (S1) represent the mechanical work done by the myosin element. The next two terms capture the increase in the cell's chemical energy associated with deviations of myosin contractility from its baseline value ($\rho_0$). In these terms, $\beta_{\mathrm{v}}$ and $\beta_{\mathrm{d}}$ denote the volumetric and deviatoric chemo-mechanical parameters, respectively, which resist the amplification of contractility in response to mechanical tension by limiting myosin phosphorylation; accordingly, higher values of $\beta_{\mathrm{v}}$ and $\beta_{\mathrm{d}}$ increase the energetic cost required to surpass the baseline contractility ($\rho_0$). Specifically, $\beta_{\mathrm{v}}$ regulates the mean level of contractility, maintaining contractility at a basal state, while $\beta_{\mathrm{d}}$ characterizes the resistance of myosin motors to align along the cell's polarization axis.

The subsequent two terms account for the reduction in the cell's chemical energy associated with myosin phosphorylation and ATP hydrolysis—processes that are promoted by mechanical tension through the activation of $Ca^{2+}$ and Rho-ROCK signaling pathways. In these terms, $\alpha_{\mathrm{v}}$ and $\alpha_{\mathrm{d}}$ represent the volumetric and deviatoric feedback parameters, respectively, which govern the strength of tension-dependent activation of contractility; as $\alpha_{\mathrm{v}}$ and $\alpha_{\mathrm{d}}$ increase, the feedback between cell contractility and tissue tension becomes more pronounced. Notably, $\alpha_{\mathrm{v}}$ modulates the overall density of myosin phosphorylation, whereas $\alpha_{\mathrm{d}}$ reflects the cell's tendency toward polarized contraction. The parameter values used in the three-dimensional model are provided in Table S1.

The final four terms represent the mechanical energy stored within the contractile unit. Since the microtubule element and myosin element are arranged in parallel, they experience the same strain, denoted by $\varepsilon_{ij}^{(\mathrm{c})}$.

Additionally, $K_{\mathrm{MT}}$ and $\mu_{\mathrm{MT}}$ denote the bulk and shear moduli of the microtubule element, which are expressed as follows:

$$K_{\mathrm{MT}} = \frac{E_{\mathrm{MT}}}{3(1-2\nu_{\mathrm{MT}})}$$
$$\mu_{\mathrm{MT}} = \frac{E_{\mathrm{MT}}}{2(1+\nu_{\mathrm{MT}})} \tag{S3}$$

where $E_{\mathrm{MT}}$ and $\nu_{\mathrm{MT}}$ are the elastic modulus and Poisson's ratio of microtubule.

Minimization of $W$ with respect to the volumetric variables $\varepsilon_{kk}^{(\mathrm{c})}$ and $\rho_{kk}$ yields the following constitutive relations:

$$\frac{\partial W}{\partial\, \varepsilon_{kk}^{(\mathrm{c})}} = \frac{1}{3}\,\rho_{kk} + K_{\mathrm{MT}}\,\varepsilon_{kk}^{(\mathrm{c})} - \frac{1}{3}\sigma_{kk} = 0 \tag{S4}$$

$$\frac{\partial W}{\partial \rho_{kk}} = \frac{1}{3}\;\; \varepsilon_{kk}^{(\mathrm{c})} + \frac{\beta_{\mathrm{v}}}{3}(\rho_{kk} - 3\rho_0) - \frac{1}{3}\alpha_{\mathrm{v}}\sigma_{kk} = 0 \tag{S5}$$

Substituting $\varepsilon_{kk}^{(\mathrm{c})}$ from (S4) into (S5) yields a constitutive relationship linking the mean myosin contractility $\frac{1}{3}\rho_{kk}$ and average stress, $\frac{1}{3}\sigma_{kk}$:

$$\frac{\rho_{kk}}{3} = \frac{3K_{\mathrm{MT}}\beta_{\mathrm{v}}}{3K_{\mathrm{MT}}\beta_{\mathrm{v}} - 1}\rho_0 + \frac{3K_{\mathrm{MT}}\alpha_{\mathrm{v}} - 1}{3K_{\mathrm{MT}}\beta_{\mathrm{v}} - 1}\frac{\sigma_{kk}}{3} \tag{S6}$$

Equation (S6) can be written in a simplified form below:

$$\frac{\rho_{kk}}{3} = f_0\rho_0 + f_m\frac{\sigma_{kk}}{3} \tag{S7}$$

As noted previously, myosin contractility is experimentally observed to increase in response to tension through the activation of $Ca^{2+}$ and Rho-Rock signaling pathways.[3–8] Equation (S7) shows how this mechanosensitive behavior is captured in our 3D model, where the average myosin contractility in all three directions, $\frac{1}{3}\rho_{kk} = (\rho_{11} + \rho_{22} + \rho_{33})/3$, is correlated with the average stress, $\frac{1}{3}\sigma_{kk} = (\sigma_{11} + \sigma_{22} + \sigma_{33})/3$. In (S7), $\rho_0$ is the baseline myosin contractility, and the constant parameter $f_0$ governs the mean contractility $\frac{1}{3}\rho_{kk}$ in the absence of stress. The constant $f_m$ serves as the regulating parameter for the tension-dependent feedback mechanism between the mean contractility and the average stress, $\frac{1}{3}\sigma_{kk}$.

Substituting $\rho_{kk}$ from (S4) into (S5) results in the following relation between $\sigma_{kk}$ and $\varepsilon_{kk}^{(\mathrm{c})}$:

$$\sigma_{kk} = \frac{3\beta_\mathrm{v}}{\beta_\mathrm{v} - \alpha_\mathrm{v}}\rho_0 + \frac{3K_\mathrm{MT}\beta_\mathrm{v} - 1}{\beta_\mathrm{v} - \alpha_\mathrm{v}}\,\varepsilon_{kk}^{(\mathrm{c})} \tag{S8}$$

Substituting back $\sigma_{kk}$ from (S8) into (S4) yields the relation below between $\rho_{kk}$ and $\varepsilon_{kk}^{(\mathrm{c})}$:

$$\rho_{kk} = \frac{3\beta_\mathrm{v}}{\beta_\mathrm{v} - \alpha_\mathrm{v}}\rho_0 + \frac{3K_\mathrm{MT}\alpha_\mathrm{v} - 1}{\beta_\mathrm{v} - \alpha_\mathrm{v}}\,\varepsilon_{kk}^{(\mathrm{c})} \tag{S9}$$

Similarly, minimization of $W$ with respect to the deviatoric variables $\tilde{\varepsilon}_{ij}^{(\mathrm{c})}$ and $\tilde{\rho}_{ij}$ yields the following relations:

$$\frac{\partial W}{\partial \tilde{\varepsilon}_{ij}^{(\mathrm{c})}} = \tilde{\rho}_{ij} + 2\mu_\mathrm{MT}\tilde{\varepsilon}_{ij}^{(\mathrm{c})} - \tilde{\sigma}_{ij} = 0 \tag{S10}$$

$$\frac{\partial W}{\partial \tilde{\rho}_{ij}} = \tilde{\varepsilon}_{ij}^{(\mathrm{c})} + \beta_\mathrm{d}\tilde{\rho}_{ij} - \alpha_\mathrm{d}\tilde{\sigma}_{ij} \tag{S11}$$

Substituting $\tilde{\sigma}_{ij}$ from (S14) into (S15) gives the following relation between $\tilde{\rho}_{ij}$ and $\tilde{\varepsilon}_{ij}^{(\mathrm{c})}$:

$$\tilde{\rho}_{ij} = \frac{2\mu_\mathrm{MT}\alpha_\mathrm{d}}{\beta_\mathrm{d} - \alpha_\mathrm{d}}\tilde{\varepsilon}_{ij}^{(\mathrm{c})} \tag{S12}$$

Substituting (S12) into (S10) results in the following relation between $\tilde{\sigma}_{ij}$ and $\tilde{\varepsilon}_{ij}^{(\mathrm{c})}$:

$$\tilde{\sigma}_{ij} = \frac{2\mu_\mathrm{MT}\beta_\mathrm{d} - 1}{\beta_\mathrm{d} - \alpha_\mathrm{d}}\tilde{\varepsilon}_{ij}^{(\mathrm{c})} \tag{S13}$$

Equations (S8), (S9), (S12), and (S13) can be written in simplified forms below:

$$\rho_{kk} = 3\bar{\rho}_0 + 3\bar{K}^{(\rho)}\varepsilon_{kk}^{(\mathrm{c})} \tag{S14}$$

$$\sigma_{kk} = 3\bar{\rho}_0 + 3\bar{K}_\mathrm{MT}\varepsilon_{kk}^{(\mathrm{c})} \tag{S15}$$

$$\tilde{\rho}_{ij} = 2\bar{\mu}^{(\rho)}\tilde{\varepsilon}_{ij}^{(\mathrm{c})} \tag{S16}$$

$$\tilde{\sigma}_{ij} = 2\bar{\mu}_{\mathrm{MT}}\tilde{\varepsilon}_{ij}^{(\mathrm{c})}\tilde{\varepsilon}_{ij}^{(\mathrm{c})} \tag{S17}$$

where the effective contractility $\bar{\rho}_0$, effective bulk modulus for microtubule $\bar{K}_{\mathrm{MT}}$, effective shear modulus for microtubule $\bar{\mu}_{\mathrm{MT}}$, effective modulus for motor density $\bar{K}_{\rho}$, and effective modulus for polarization $\bar{\mu}_{\rho}$ are defined as:

$$\begin{aligned}
\bar{\rho}_0 &= \frac{\beta_{\mathrm{v}}}{\beta_{\mathrm{v}} - \alpha_{\mathrm{v}}}\rho_0 \\
\bar{K}_{\mathrm{MT}} &= \frac{3K_{\mathrm{MT}}\beta_{\mathrm{v}} - 1}{3(\beta_{\mathrm{v}} - \alpha_{\mathrm{v}})} \\
\bar{K}_{\rho} &= \frac{3K_{\mathrm{MT}}\alpha_{\mathrm{v}} - 1}{3(\beta_{\mathrm{v}} - \alpha_{\mathrm{v}})} \\
\bar{\mu}_{\mathrm{MT}} &= \frac{2\mu_{\mathrm{MT}}\beta_{\mathrm{d}} - 1}{2(\beta_{\mathrm{d}} - \alpha_{\mathrm{d}})} \\
\bar{\mu}_{\rho} &= \frac{2\mu_{\mathrm{MT}}\alpha_{\mathrm{d}} - 1}{2(\beta_{\mathrm{d}} - \alpha_{\mathrm{d}})}
\end{aligned} \tag{S18}$$

To implement the model within a three-dimensional finite element framework, we need to derive expressions for the total stress tensor $\sigma_{ij}$ and the total stiffness tensor $C_{ij}$. For this purpose, we need to derive the expression for contractility tensor $\rho_{kk}$. To derive the contractility tensor, we substitute $\rho_{kk}$ and $\tilde{\rho}_{ij}$ from (S14) and (S16) into (S2) where $\rho_{ij} = \frac{1}{3}\rho_{kk}\delta_{ij} + \tilde{\rho}_{ij}$, which yields the following expression:

$$\rho_{ij} = \bar{K}_{\rho}\varepsilon_{kk}^{(\mathrm{c})}\delta_{ij} + 2\bar{\mu}_{\rho}\left(\varepsilon_{ij}^{(\mathrm{c})} - \frac{1}{3}\varepsilon_{kk}^{(\mathrm{c})}\delta_{ij}\right) + \bar{\rho}_0\delta_{ij} \tag{S19}$$

Part of the contractility $\rho_{ij}$ compressively loads the microtubule network, inducing compressive stress $C_{ijkl}^{(\mathrm{MT})}\varepsilon_{kl}^{(\mathrm{c})}$, where the fourth order tensor $C_{ijkl}^{(\mathrm{MT})}$ is the stiffness of microtubule expressed as follows:

$$C_{ijkl}^{(\mathrm{MT})} = K_{\mathrm{MT}}\delta_{ij}\delta_{kl} + \mu_{\mathrm{MT}}(\delta_{ik}\delta_{jk} + \delta_{il}\delta_{jk} - \frac{2}{3}\delta_{ij}\delta_{kl}) \tag{S20}$$

Similarly, the rest of the cell contractility is transmitted as stress $\sigma_{ij}$, generating tensile stress in the cytoskeleton:

$$\rho_{ij} = -C_{ijkl}^{(\mathrm{MT})}\varepsilon_{kl}^{(\mathrm{c})} + \sigma_{ij} \tag{S21}$$

In our model, the contractility tensor $\rho_{ij}$ is initially isotropic, exhibiting equal contractility in all directions in the initial configuration. This initial isotropic contractility can be mathematically expressed by rewriting (S19) in the following form:

$$\rho_{ij} = C_{ijkl}^{(\rho)}\varepsilon_{kl}^{(\mathrm{c})} + \bar{\rho}_0\delta_{ij} \tag{S22}$$

where $C_{ijkl}^{(\rho)}$ is a fourth-order myosin stiffness tensor,

$$C_{ijkl}^{(\rho)} = \bar{K}_{\rho}\delta_{ij}\delta_{kl} + \bar{\mu}_{\rho}(\delta_{ik}\delta_{jk} + \delta_{il}\delta_{jk} - \frac{2}{3}\delta_{ij}\delta_{kl}) \tag{S23}$$

Equations (S21) and (S22) demonstrate that the contractility tensor $\rho_{ij}$ is isotropic under stress-free conditions ($\sigma_{ij} = 0$), where the diagonal components of $\rho_{ij}$ are all equal and non-zero $\rho_{11} = \rho_{22} = \rho_{33} \neq 0$, while all off-diagonal components are zero.

Cells also respond to mechanical tension by polymerizing actin filaments and organizing them into aligned bundles oriented along the direction of applied tension.[9,10] To capture the strain-stiffening behavior of actin network, we hypothesize that the stiffness of the actin network increases proportionally with, and aligned along, the tensile principal components of the stress tensor $\sigma_{ij}$:

$$C_{ijkl}^{(A)} = C_{ijkl}^{(A0)} + C_{ijkl}^{(AS)} \tag{S24}$$

where $C_{ijkl}^{(A0)}$ is the initial stiffness of the actin network, and $C_{ijkl}^{(AS)}$ represents the stiffening of actin under tension. The initial stiffness tensor of actin $C_{ijkl}^{(A0)}$ can be written as:

$$C_{ijkl}^{(A0)} = K_{A0}\delta_{ij}\delta_{kl} + \mu_{A0}(\delta_{ik}\delta_{jk} + \delta_{il}\delta_{jk} - \frac{2}{3}\delta_{ij}\delta_{kl}) \tag{S25}$$

where $K_{A0}$ and $\mu_{A0}$ denote the initial bulk and the shear moduli of actin, and are expressed as follows:

$$K_{A0} = \frac{E_{A0}}{3(1-2\nu_{A0})}$$
$$\mu_{A0} = \frac{E_{A0}}{2(1+\nu_{A0})} \tag{S26}$$

To define $C_{ijkl}^{(AS)}$, we decompose the tensile stress $\sigma_{ij}$ in the following manner:

$$\sigma_{ij} = \sigma_{ij}^{(A)} = \sigma_{ij}^{(A0)} + \sigma_{ij}^{(AS)} \tag{S27}$$

where $\sigma^{(A0)}$ is linearly linked to the strain in tensile element $\varepsilon^{(t)}$, through the initial stiffness tensor:

$$\sigma_{ij}^{(A0)} = C_{ijkl}^{(A0)}\varepsilon_{kl}^{(t)} \tag{S28}$$

where $\varepsilon_{kl}^{(t)}$ is the three-dimensional representation $\varepsilon^{(t)}$ in Fig. S7. We then determine the eigenvalues (principal strains) and the eigenprojections ($E_1 = n_1 \otimes n_1$, $E_2 = n_2 \otimes n_2$, $E_3 = n_3 \otimes n_3$) of the strain tensor $\varepsilon^{(t)}$:

$$\varepsilon^{(t)} = \sum_{i=1}^{3} \varepsilon_i^{(t)}\, n_i \otimes n_i = \sum_{i=1}^{3} \varepsilon_i^{(t)}\, E_i \tag{S29}$$

Where eigenvectors $n_1$, $n_2$, and $n_3$ are the orthogonal unit vectors in the direction of eigenvalues $\varepsilon_1^{(t)}$, $\varepsilon_2^{(t)}$, and $\varepsilon_3^{(t)}$, respectively. In (S29), $\otimes$ represents the dyadic product of two vectors which is defined as $(u \otimes v)_{ij} = u_i v_j$. Using the eigenvalues and the eigenprojections of $\varepsilon^{(\mathrm{t})}$, we define the stress tensor $\sigma_{ij}^{(\mathrm{AS})}$ as a nonlinear function of the strain tensor:

$$\sigma^{(\mathrm{AS})}=\sum_{i=1}^{3}\frac{\partial f(\varepsilon_i^{(\mathrm{t})})}{\partial \varepsilon_i^{(\mathrm{t})}}n_i\otimes n_i=\sum_{i=1}^{3}\sigma^{(\mathrm{AS})}\left(\varepsilon_i^{(\mathrm{t})}\right)E_i=\sum_{i=1}^{3}\sigma_i^{(\mathrm{AS})}E_i \tag{S30}$$

where $f(\varepsilon_i^{(\mathrm{t})})$ is the energy function and its derivative is defined in the following form to ensure the continuity of the first and second derivatives of $\sigma_i^{(\mathrm{AS})}$ with respect to $\varepsilon_i^{(\mathrm{t})}$:

$$\sigma_i^{(\mathrm{AS})}=\frac{\partial f(\varepsilon_i^{(\mathrm{t})})}{\partial \varepsilon_i^{(\mathrm{t})}}$$
$$=\begin{cases}0 & \varepsilon_i^{(\mathrm{t})}<\epsilon_1\\ \ell\dfrac{\left(\dfrac{\varepsilon_i^{(\mathrm{t})}-\epsilon_1}{\epsilon_2-\epsilon_1}\right)^n\left(\varepsilon_i^{(\mathrm{t})}-\epsilon_1\right)^2}{(n+1)(n+2)} & \epsilon_1\le\varepsilon_i^{(\mathrm{t})}<\epsilon_2\\ \ell\left[\dfrac{\left(1+\varepsilon_i^{(\mathrm{t})}-\epsilon_2\right)^{m+2}-1}{(m+1)(m+2)}+\dfrac{\epsilon_2-\varepsilon_i^{(\mathrm{t})}}{m+1}+\dfrac{(\varepsilon_i^{(\mathrm{t})}-\epsilon_2)(\epsilon_2-\epsilon_1)}{n+1}+\dfrac{(\epsilon_2-\epsilon_1)^2}{(n+1)(n+2)}\right] & \varepsilon_i^{(\mathrm{t})}\ge\epsilon_2\end{cases} \tag{S31}$$

In (S31), $\epsilon_1=\epsilon_{\mathrm{c}}-0.5\epsilon_{\mathrm{t}}$ and $\epsilon_2=\epsilon_{\mathrm{c}}+0.5\epsilon_{\mathrm{t}}$, where $\epsilon_{\mathrm{c}}$ and $\epsilon_{\mathrm{t}}=0.25\epsilon_{\mathrm{c}}$ are the critical principal strain and transition width, respectively. For small strains, strain-stiffening is zero ($\sigma_i^{(\mathrm{ts})}=0$), while for larger strains, the principal stress $\sigma_i^{(\mathrm{ts})}$ rises nonlinearly with the principal strain $\varepsilon_i^{(\mathrm{t})}$ with a transition region between $\epsilon_1$ and $\epsilon_2$, where $\ell$ and $m$ are strain-stiffening parameters that regulate the rise in stiffness, and $n$ is the transition constant. With $\varepsilon^{(\mathrm{t})}$ and $\sigma^{(\mathrm{ts})}$ at hand from (S29) and (S30), $C^{(\mathrm{AS})}$ in (S24) can be determined using the following linear approximation:

$$C_{ijkl}^{(\mathrm{AS})}=\frac{d\sigma_{ij}^{(\mathrm{AS})}}{d\varepsilon_{kl}^{(\mathrm{t})}}\qquad\text{or}\qquad C^{(\mathrm{AS})}=\frac{d\sigma^{(\mathrm{AS})}}{d\varepsilon^{(\mathrm{t})}} \tag{S32}$$

Note that (S31) provides $\sigma_i^{(\mathrm{AS})}$ (rather than $\sigma_{ij}^{(\mathrm{AS})}$) as a function of $\varepsilon_i^{(\mathrm{t})}$. We first use the definition of spectral decomposition of $\sigma^{(\mathrm{AS})}$ in (S30) to rewrite (S32) in the following form:

$$C^{(\mathrm{AS})}=\sum_{i=1}^{3}(E_i\otimes\frac{d\sigma_i^{(\mathrm{AS})}}{d\varepsilon^{(\mathrm{t})}}+\sigma_i^{(\mathrm{AS})}\frac{dE_i}{d\varepsilon^{(\mathrm{t})}}) \tag{S33}$$

We then apply the chain rule to evaluate the first term in (S33)

$$C^{(\mathrm{AS})}=\sum_{i=1}^{3}(\sum_{j=1}^{3}\frac{\partial\sigma_i^{(\mathrm{AS})}}{\partial\varepsilon_j^{(\mathrm{t})}}E_i\otimes\frac{d\varepsilon_j^{(\mathrm{t})}}{d\varepsilon^{(\mathrm{t})}}+\sigma_i^{(\mathrm{AS})}\frac{dE_i}{d\varepsilon^{(\mathrm{t})}}) \tag{S34}$$

To derive an exact expression for $C^{(\mathrm{AS})}$, we consider three possible cases based on the number of distinct eigenvalues for $\varepsilon_{ij}^{(\mathrm{t})}$. In the first case, where all three eigenvalues of the strain tensor $\varepsilon_{ij}^{(\mathrm{t})}$ are nonidentical ($\varepsilon_1^{(\mathrm{t})} \neq \varepsilon_2^{(\mathrm{t})} \neq \varepsilon_3^{(\mathrm{t})}$), we can determine the following expression for $C^{(\mathrm{AS})}$ from (S34) by taking derivatives of $\varepsilon_j^{(\mathrm{t})}$ and $E_i$ with respect to $\varepsilon^{(\mathrm{t})}$:

$$\begin{aligned} C^{(AS)} = \sum_{a=1}^{3} \frac{\sigma_a^{(\mathrm{AS})}}{(\varepsilon_a^{(\mathrm{t})} - \varepsilon_b^{(\mathrm{t})})(\varepsilon_a^{(\mathrm{t})} - \varepsilon_c^{(\mathrm{t})})} \Bigg\{ & \frac{d\left(\varepsilon^{(\mathrm{t})}\right)^2}{d\varepsilon^{(\mathrm{t})}} - \left(\varepsilon_b^{(\mathrm{t})} + \varepsilon_c^{(\mathrm{t})}\right) I_\mathrm{s} \\ & - \left[\left(\varepsilon_a^{(\mathrm{t})} - \varepsilon_b^{(\mathrm{t})}\right) + \left(\varepsilon_a^{(\mathrm{t})} - \varepsilon_c^{(\mathrm{t})}\right)\right] E_a \otimes E_a - \left(\varepsilon_b^{(\mathrm{t})} - \varepsilon_c^{(\mathrm{t})}\right)(E_b \otimes E_b - E_c \otimes E_c) \Bigg\} \\ & + \sum_{i=1}^{3} \sum_{j=1}^{3} \frac{\partial \sigma_i^{(\mathrm{AS})}}{\partial \varepsilon_j^{(\mathrm{t})}} E_i \otimes E_j \end{aligned} \qquad \text{(S35)}$$

where $(a, b, c)$ are cyclic permutations of (1, 2, 3), $I_\mathrm{s}$ is the fourth-order symmetric identity tensor,

$$(I_\mathrm{s})_{ijkl} = \frac{1}{2}\left(I + I^\mathrm{T}\right) = \frac{1}{2}\left(\delta_{ik}\delta_{jl} + \delta_{il}\delta_{jk}\right) \qquad \text{(S36)}$$

and $\frac{d\left(\varepsilon^{(\mathrm{t})}\right)^2}{d\varepsilon^{(\mathrm{t})}}$ is a fourth-order tensor and derivative of the square of the second order tensor $\varepsilon^{(\mathrm{t})}$:

$$\left(\frac{d\left(\varepsilon^{(\mathrm{t})}\right)^2}{d\varepsilon^{(\mathrm{t})}}\right)_{ijkl} = \frac{1}{2}\left(\delta_{ik}\varepsilon_{lj}^{(\mathrm{t})} + \delta_{il}\varepsilon_{kj}^{(\mathrm{t})} + \delta_{jl}\varepsilon_{ik}^{(\mathrm{t})} + \delta_{kj}\varepsilon_{il}^{(\mathrm{t})}\right) \qquad \text{(S37)}$$

In the second case, $\varepsilon_{ij}^{(\mathrm{t})}$ has two identical eigenvalues ($\varepsilon_1^{(\mathrm{t})} \neq \varepsilon_2^{(\mathrm{t})} = \varepsilon_3^{(\mathrm{t})}$) which gives the following expression for $C^{(\mathrm{AS})}$:

$$C^{(\mathrm{AS})} = s_1 \frac{d\left(\varepsilon^{(\mathrm{t})}\right)^2}{d\varepsilon^{(\mathrm{t})}} - s_2 I_\mathrm{s} - s_3 \varepsilon^{(\mathrm{t})} \otimes \varepsilon^{(\mathrm{t})} + s_4 \varepsilon^{(\mathrm{t})} \otimes I + s_5 I \otimes \varepsilon^{(\mathrm{t})} - s_6 I \otimes I \qquad \text{(S38)}$$

where $I_{ij} = \delta_{ij}$ is the second-order identity tensor, and constants $s_i$ are as follows:

$$\begin{aligned} s_1 &= \frac{\sigma_a^{(\mathrm{AS})} - \sigma_c^{(\mathrm{AS})}}{\left(\varepsilon_a^{(\mathrm{t})} - \varepsilon_c^{(\mathrm{t})}\right)^2} + \frac{1}{\varepsilon_a^{(\mathrm{t})} - \varepsilon_c^{(\mathrm{t})}}\left(\frac{\partial \sigma_c^{(\mathrm{AS})}}{\partial \varepsilon_b^{(\mathrm{t})}} - \frac{\partial \sigma_c^{(\mathrm{AS})}}{\partial \varepsilon_c^{(\mathrm{t})}}\right) \\ s_2 &= 2\varepsilon_c^{(\mathrm{t})} \frac{\sigma_a^{(\mathrm{AS})} - \sigma_c^{(\mathrm{AS})}}{\left(\varepsilon_a^{(\mathrm{t})} - \varepsilon_c^{(\mathrm{t})}\right)^2} + \frac{\varepsilon_a^{(\mathrm{t})} + \varepsilon_c^{(\mathrm{t})}}{\varepsilon_a^{(\mathrm{t})} - \varepsilon_c^{(\mathrm{t})}}\left(\frac{\partial \sigma_c^{(\mathrm{AS})}}{\partial \varepsilon_b^{(\mathrm{t})}} - \frac{\partial \sigma_c^{(\mathrm{AS})}}{\partial \varepsilon_c^{(\mathrm{t})}}\right) \\ s_3 &= 2\frac{\sigma_a^{(\mathrm{AS})} - \sigma_c^{(\mathrm{AS})}}{\left(\varepsilon_a^{(\mathrm{t})} - \varepsilon_c^{(\mathrm{t})}\right)^3} + \frac{1}{\left(\varepsilon_a^{(\mathrm{t})} - \varepsilon_c^{(\mathrm{t})}\right)^2}\left(\frac{\partial \sigma_a^{(\mathrm{AS})}}{\partial \varepsilon_c^{(\mathrm{t})}} + \frac{\partial \sigma_c^{(\mathrm{AS})}}{\partial \varepsilon_a^{(\mathrm{t})}} - \frac{\partial \sigma_a^{(\mathrm{AS})}}{\partial \varepsilon_a^{(\mathrm{t})}} - \frac{\partial \sigma_c^{(\mathrm{AS})}}{\partial \varepsilon_c^{(\mathrm{t})}}\right) \end{aligned} \qquad \text{(S39)}$$

$$s_4 = 2\varepsilon_c^{(\mathrm{t})}\frac{\sigma_a^{(\mathrm{AS})} - \sigma_c^{(\mathrm{AS})}}{\left(\varepsilon_a^{(\mathrm{t})} - \varepsilon_c^{(\mathrm{t})}\right)^3} + \frac{1}{\varepsilon_a^{(\mathrm{t})} - \varepsilon_c^{(\mathrm{t})}}\left(\frac{\partial\sigma_a^{(\mathrm{AS})}}{\partial\varepsilon_c^{(\mathrm{t})}} - \frac{\partial\sigma_c^{(\mathrm{AS})}}{\partial\varepsilon_b^{(\mathrm{t})}}\right) + \frac{\varepsilon_c^{(\mathrm{t})}}{\left(\varepsilon_a^{(\mathrm{t})} - \varepsilon_c^{(\mathrm{t})}\right)^2}\left(\frac{\partial\sigma_a^{(\mathrm{AS})}}{\partial\varepsilon_c^{(\mathrm{t})}} + \frac{\partial\sigma_c^{(\mathrm{AS})}}{\partial\varepsilon_a^{(\mathrm{t})}} - \frac{\partial\sigma_a^{(\mathrm{AS})}}{\partial\varepsilon_a^{(\mathrm{t})}} - \frac{\partial\sigma_c^{(\mathrm{AS})}}{\partial\varepsilon_c^{(\mathrm{t})}}\right)$$

$$s_5 = 2\varepsilon_c^{(\mathrm{t})}\frac{\sigma_a^{(\mathrm{AS})} - \sigma_c^{(\mathrm{AS})}}{\left(\varepsilon_a^{(\mathrm{t})} - \varepsilon_c^{(\mathrm{t})}\right)^3} + \frac{1}{\varepsilon_a^{(\mathrm{t})} - \varepsilon_c^{(\mathrm{t})}}\left(\frac{\partial\sigma_c^{(\mathrm{AS})}}{\partial\varepsilon_a^{(\mathrm{t})}} - \frac{\partial\sigma_c^{(\mathrm{AS})}}{\partial\varepsilon_b^{(\mathrm{t})}}\right) + \frac{\varepsilon_c^{(\mathrm{t})}}{\left(\varepsilon_a^{(\mathrm{t})} - \varepsilon_c^{(\mathrm{t})}\right)^2}\left(\frac{\partial\sigma_a^{(\mathrm{AS})}}{\partial\varepsilon_c^{(\mathrm{t})}} + \frac{\partial\sigma_c^{(\mathrm{AS})}}{\partial\varepsilon_a^{(\mathrm{t})}} - \frac{\partial\sigma_a^{(\mathrm{AS})}}{\partial\varepsilon_a^{(\mathrm{t})}} - \frac{\partial\sigma_c^{(\mathrm{AS})}}{\partial\varepsilon_c^{(\mathrm{t})}}\right)$$

$$s_6 = 2\varepsilon_c^{(\mathrm{t})}\frac{\sigma_a^{(\mathrm{AS})} - \sigma_c^{(\mathrm{AS})}}{\left(\varepsilon_a^{(\mathrm{t})} - \varepsilon_c^{(\mathrm{t})}\right)^3} + \frac{\varepsilon_a^{(\mathrm{AS})}\varepsilon_c^{(\mathrm{AS})}}{\left(\varepsilon_a^{(\mathrm{t})} - \varepsilon_c^{(\mathrm{t})}\right)^2}\left(\frac{\partial\sigma_a^{(\mathrm{AS})}}{\partial\varepsilon_c^{(\mathrm{t})}} + \frac{\partial\sigma_c^{(\mathrm{AS})}}{\partial\varepsilon_a^{(\mathrm{t})}}\right) - \frac{\left(\varepsilon_c^{(\mathrm{t})}\right)^2}{\left(\varepsilon_a^{(\mathrm{t})} - \varepsilon_c^{(\mathrm{t})}\right)^2}\left(\frac{\partial\sigma_a^{(\mathrm{AS})}}{\partial\varepsilon_a^{(\mathrm{t})}} + \frac{\partial\sigma_c^{(\mathrm{AS})}}{\partial\varepsilon_c^{(\mathrm{t})}}\right) - \frac{\varepsilon_a^{(\mathrm{t})} + \varepsilon_c^{(\mathrm{t})}}{\varepsilon_a^{(\mathrm{t})} - \varepsilon_c^{(\mathrm{t})}}\frac{\partial\sigma_c^{(\mathrm{AS})}}{\partial\varepsilon_b^{(\mathrm{t})}}$$

In the third case, all eigenvalues are identical ($\varepsilon_1^{(\mathrm{t})} = \varepsilon_2^{(\mathrm{t})} = \varepsilon_3^{(\mathrm{t})}$), which gives the following expression for $C^{(\mathrm{ts})}$:

$$C^{(\mathrm{AS})} = \left(\frac{\partial\sigma_1^{(\mathrm{AS})}}{\partial\varepsilon_1^{(\mathrm{t})}} - \frac{\partial\sigma_1^{(\mathrm{AS})}}{\partial\varepsilon_2^{(\mathrm{t})}}\right)I_{\mathrm{s}} + \frac{\partial\sigma_1^{(\mathrm{AS})}}{\partial\varepsilon_2^{(\mathrm{t})}} I \otimes I \tag{S40}$$

Note that we require $\partial\sigma_i^{(\mathrm{AS})}/\partial\varepsilon_j^{(\mathrm{t})}$ for all three cases, and this can be determined by calculating the first derivative of $\sigma_i^{(\mathrm{AS})}$ in (S32).

$$\frac{\partial\sigma_i^{(\mathrm{AS})}}{\partial\varepsilon_i^{(\mathrm{t})}} = \frac{\partial}{\partial\varepsilon_i^{(\mathrm{t})}}\left(\frac{\partial f}{\partial\varepsilon_i^{(\mathrm{t})}}\right) = \begin{cases} 0, & \varepsilon_i^{(\mathrm{t})} < \epsilon_1 \\ \ell\dfrac{\left(\dfrac{\varepsilon_i^{(\mathrm{t})} - \epsilon_1}{\epsilon_2 - \epsilon_1}\right)^n \left(\varepsilon_i^{(\mathrm{t})} - \epsilon_1\right)}{n+1}, & \epsilon_1 \le \varepsilon_i^{(\mathrm{t})} < \epsilon_2 \\ \ell\left[\dfrac{\left(1 + \varepsilon_i^{(\mathrm{t})} - \epsilon_2\right)^{m+1} - 1}{m+1} + \dfrac{\epsilon_2 - \epsilon_1}{n+1}\right], & \varepsilon_i^{(\mathrm{t})} \ge \epsilon_2 \end{cases} \tag{S41}$$

Note that $\sigma_i^{(\mathrm{AS})}$ in (S41) is only a function of $\varepsilon_i^{(\mathrm{t})}$ and is independent of $\varepsilon_j^{(\mathrm{t})}$ for $i \neq j$. Having $C^{(\mathrm{AS})}$, the stiffness of actin $C^{(\mathrm{A})}$ can be obtained from (S24).

The cell model, as previously described, consists of three elements: myosin with the stiffness tensor $C_{ijkl}^{(\rho)}$, microtubule with the stiffness tensor $C_{ijkl}^{(\mathrm{MT})}$, and actin with the stiffness tensor $C_{ijkl}^{(\mathrm{A})}$. To obtain the total stiffness of the cell, first degrade the fourth-order stiffness tensors to second-order stiffness tensors using (S42).

$$C_{ij} = \begin{bmatrix} C_{1111} & C_{1122} & C_{1133} & C_{1112} & C_{1113} & C_{1123} \\ C_{2211} & C_{2222} & C_{2233} & C_{2212} & C_{2213} & C_{2223} \\ C_{3311} & C_{3322} & C_{3333} & C_{3312} & C_{3313} & C_{3323} \\ C_{1211} & C_{1222} & C_{1233} & C_{1212} & C_{1213} & C_{1223} \\ C_{1311} & C_{1322} & C_{1333} & C_{1312} & C_{1313} & C_{1323} \\ C_{2311} & C_{2322} & C_{2333} & C_{2312} & C_{2313} & C_{2323} \end{bmatrix} \tag{S42}$$

Given that the myosin and microtubule elements are connected in parallel, and this combined unit is linked in series with the actin element, the total stiffness tensor of the cell, $C$ is obtained as follows:

$$C = \left(\left(C^{(\rho)} + C^{(\mathrm{MT})}\right)^{-1} + (C^{(\mathrm{A})})^{-1}\right)^{-1} \tag{S43}$$

As described in the derived equations, both the stress tensor $\sigma_{ij}$ (from (S21) and (S27)) and stiffness tensor $C_{ij}$ (from (S43)) are functions of unknown strain tensors $\varepsilon_{ij}^{(\mathrm{c})}$ and $\varepsilon_{ij}^{(\mathrm{t})}$. Therefore, we use an iterative procedure to determine the unknowns $\varepsilon_{ij}^{(\mathrm{c})}$ and $\varepsilon_{ij}^{(\mathrm{t})}$. We first define the following 12×1 vector which contains all 12 unknown variables in the strain tensors $\varepsilon_{ij}^{(\mathrm{c})}$ and $\varepsilon_{ij}^{(\mathrm{t})}$.

$$\begin{aligned} u &= \{u_1 \quad u_2 \quad \cdots \quad u_{12}\}^{\mathrm{T}} \\ &= \left\{\varepsilon_{11}^{(\mathrm{c})} \quad \varepsilon_{22}^{(\mathrm{c})} \quad \varepsilon_{33}^{(\mathrm{c})} \quad \varepsilon_{12}^{(\mathrm{c})} \quad \varepsilon_{13}^{(\mathrm{c})} \quad \varepsilon_{23}^{(\mathrm{c})} \quad \varepsilon_{11}^{(\mathrm{t})} \quad \varepsilon_{22}^{(\mathrm{t})} \quad \varepsilon_{33}^{(\mathrm{t})} \quad \varepsilon_{12}^{(\mathrm{t})} \quad \varepsilon_{13}^{(\mathrm{t})} \quad \varepsilon_{23}^{(\mathrm{t})}\right\}^{\mathrm{T}} \end{aligned} \tag{S44}$$

We need 12 equations to determine the 12 unknowns. As actin is connected in series to the other two element and the cytoskeletal stress is transmitted to this element, we use the following condition which gives 6 equations:

$$\sigma = \sigma^{(\mathrm{c})} = \sigma^{(\mathrm{A})} \tag{S45}$$

where $\sigma^{(\mathrm{c})}$ is the stress generated in the contractile section and is directly transmitted to the tissue and is given by (S21) and (S22) as follows:

$$\sigma_{ij}^{(\mathrm{c})} = \sigma_{ij} = \bar{\rho}_0 \delta_{ij} + \left(C_{ijkl}^{(\rho)} + C_{ijkl}^{(\mathrm{MT})}\right) \varepsilon_{kl}^{(\mathrm{c})} \tag{S46}$$

and $\sigma^{(\mathrm{A})}$ is the stress in the actin element which is transmitted through the cytoskeleton and is given by (S27). The other 6 equations can be obtained from the following condition:

$$\varepsilon = \varepsilon^{(\mathrm{c})} + \varepsilon^{(\mathrm{t})} \tag{S47}$$

where $\varepsilon$ is the second-order tensor of total cell strain. Note that all stress and strain tensors $\sigma_{ij}^{(\mathrm{c})}$, $\sigma_{ij}^{(\mathrm{A})}$, $\varepsilon_{ij}^{(\mathrm{c})}$, and $\varepsilon_{ij}^{(\mathrm{t})}$ are symmetric. Therefore, the 12 equations from (S45) and (S47) can be defined by the following 12 equations in the 12×1 vector $f$:

$$f = \{f_1 \quad f_2 \quad \cdots \quad f_{12}\}^{\mathrm{T}} \tag{S48}$$

where

$$
\begin{aligned}
f_1 &= \sigma_{11}^{(c)} - \sigma_{11}^{(t)} \\
f_2 &= \sigma_{22}^{(c)} - \sigma_{22}^{(t)} \\
f_3 &= \sigma_{33}^{(c)} - \sigma_{33}^{(t)} \\
f_4 &= \sigma_{12}^{(c)} - \sigma_{12}^{(t)} \\
f_5 &= \sigma_{13}^{(c)} - \sigma_{13}^{(t)} \\
f_6 &= \sigma_{23}^{(c)} - \sigma_{23}^{(t)} \\
f_7 &= \varepsilon_{11} - \varepsilon_{11}^{(c)} - \varepsilon_{11}^{(t)} \\
f_8 &= \varepsilon_{22} - \varepsilon_{22}^{(c)} - \varepsilon_{22}^{(t)} \\
f_9 &= \varepsilon_{33} - \varepsilon_{33}^{(c)} - \varepsilon_{33}^{(t)} \\
f_{10} &= \varepsilon_{12} - \varepsilon_{12}^{(c)} - \varepsilon_{12}^{(t)} \\
f_{11} &= \varepsilon_{13} - \varepsilon_{13}^{(c)} - \varepsilon_{13}^{(t)} \\
f_{12} &= \varepsilon_{23} - \varepsilon_{23}^{(c)} - \varepsilon_{23}^{(t)}
\end{aligned}
\qquad \text{(S49)}
$$

The 12×12 Jacobian matrix is then determined using (S44) and (S48).

$$
J = \begin{bmatrix} \partial f_1/\partial u_1 & \partial f_1/\partial u_2 & \dots & \partial f_1/\partial u_{12} \\ \partial f_2/\partial u_1 & \partial f_2/\partial u_2 & \dots & \partial f_2/\partial u_{12} \\ \vdots & \vdots & & \vdots \\ \partial f_{12}/\partial u_1 & \partial f_{12}/\partial u_2 & \dots & \partial f_{12}/\partial u_{12} \end{bmatrix} = \begin{bmatrix} C^{(\rho)} + C^{(c)} & -C^{(t)} \\ -I & -I \end{bmatrix} \qquad \text{(S50)}
$$

Finally, we implement the Newton-Raphson method to find the unknown vector $u$.

$$
u_{i+1} = u_i - J^{-1} f(u_i) \qquad \text{(S51)}
$$

where $u_i$ and $u_{i+1}$ are the solutions in iterations $i$ and $i+1$, respectively. The initial guess $u_0$ is:

$$
u_0 = \{0 \quad 0 \quad \dots \quad 0\}_{1\times 12}^{T} \qquad \text{(S52)}
$$

and the convergence criterion for solving the equations is:

$$
|f| = \sqrt{(f_1)^2 + (f_2)^2 + \cdots + (f_{12})^2} < \epsilon_{\mathrm{Tol}} \qquad \text{(S53)}
$$

where $|f|$ denotes the magnitude of the vector $f$, and $\epsilon_{\mathrm{Tol}}$ is the prescribed convergence threshold. In our simulation, we set $\epsilon_{Tol} = 10^{-8}$, such that the iterative procedure defined in (S51) is terminated once $|f| < 10^{-8}$. Having $\varepsilon_{ij}^{(c)}$ and $\varepsilon_{ij}^{(t)}$ obtained from (S51), we can calculate the total stiffness of the cell $C_{ij}$ from (S43) and the stress field $\sigma_{ij}$ from (S46).

Note that, the parameters $\bar{\rho}_0$, $\bar{K}_{\mathrm{MT}}$, $\bar{K}_{\rho}$, $\bar{\mu}_{\mathrm{MT}}$, and $\bar{\mu}_{\rho}$ in (S18) must all be positive to ensure chemical and mechanical stability. This leads to the following stability criterion for the model:

$$\frac{1}{3K_{\mathrm{MT}}} < \alpha_{\mathrm{v}} < \beta_{\mathrm{v}}$$
$$\frac{1}{2\mu_{\mathrm{MT}}} < \alpha_{\mathrm{d}} < \beta_{\mathrm{d}} \tag{S54}$$

We have previously shown that the feedback mechanism between mean contractility $\rho_{kk}$ and mean stress $\sigma_{kk}$, as described in (S6), predicts an increase in cell contractility, cytoskeletal tension, and traction forces with increasing substrate stiffness and cell spreading area—trends that are consistent with experimental findings.[11] However, we have recently found that, in addition to the magnitude of tension, cells also respond to its directional anisotropy.[2]

To account for this, we extended the model to incorporate the effect of tension anisotropy on myosin phosphorylation. Specifically, an additional term $\alpha_{\mathrm{a}}\sigma_{\mathrm{a}}$ was introduced to capture the enhancement of myosin phosphorylation in response to anisotropic stress distributions. The modified expression for mean contractility becomes:

$$\frac{\rho_{kk}}{3} = \frac{3K_{\mathrm{MT}}\beta_{\mathrm{v}}}{3K_{\mathrm{MT}}\beta_{\mathrm{v}} - 1}\rho_0 + \alpha_{\mathrm{a}}\sigma_{\mathrm{a}} + \frac{3K_{\mathrm{MT}}\alpha_{\mathrm{v}} - 1}{3K_{\mathrm{MT}}\beta_{\mathrm{v}} - 1}\frac{\sigma_{kk}}{3} \tag{S55}$$

where $\sigma_1 > \sigma_2 > \sigma_3$ are the principal stress components (eigenvalues) of the stress tensor $\sigma_{ij}$ with $\sigma_1 > \sigma_2 > 0$, $\sigma_{\mathrm{a}} = \tanh\left(\frac{1}{2}\left(\frac{\sigma_1}{\sigma_2} - 1\right)\right)\rho_0$ represents the tension anisotropy, $\alpha_{\mathrm{a}}$ is the anisotropic chemo-mechanical feedback parameter which regulates the enhancement in myosin phosphorylation in response to tension anisotropy. The additional term $\alpha_{\mathrm{a}}\sigma_{\mathrm{a}}$ is implemented into the finite element framework using a piecewise linear approximation where $\rho_0$ in the expression for the effective contractility $\bar{\rho}_0$ in (S18) is simply replaced with $\rho_0 + \left(\frac{3K^{(\mathrm{MT})}\beta_{\mathrm{v}} - 1}{3K^{(\mathrm{MT})}\beta_{\mathrm{v}}}\right)\alpha_{\mathrm{a}}\sigma_{\mathrm{a}}$ in each step of the simulation.

### Supplementary Figures

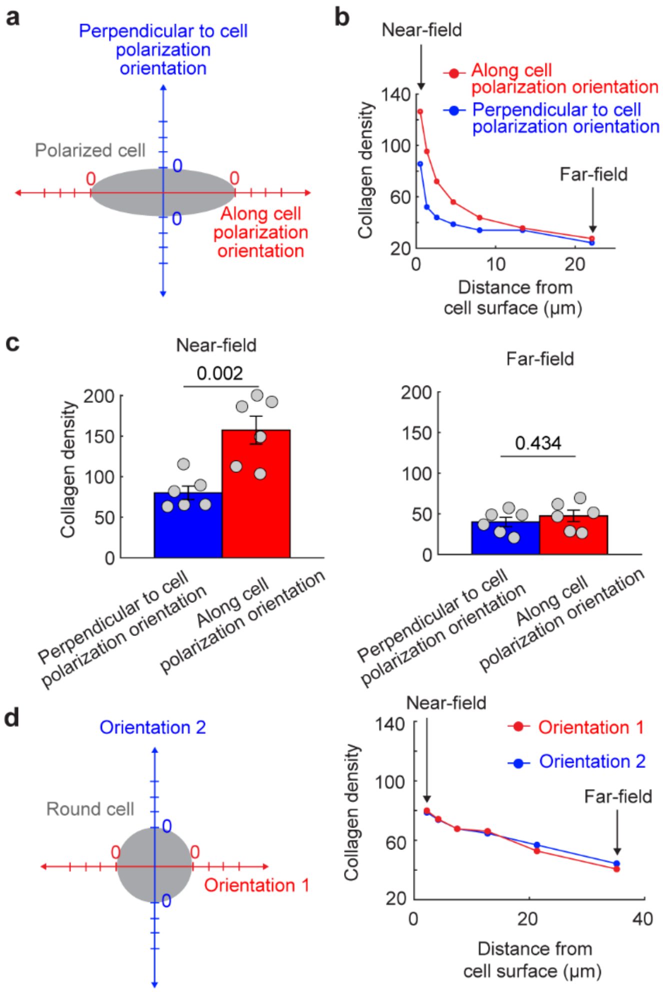


**Figure S1. Polarized fibroblasts induce anisotropic ECM remodeling. a**, We quantified collagen density along and perpendicular to the cell's polarization orientation. **b-c**, Quantitative analysis revealed that in regions distant from the cell (far-field), collagen density was comparable across orientations. However, in regions adjacent to the cell (near-field), collagen density was significantly higher along the cell's polarization orientation compared to the perpendicular direction, suggesting preferential matrix densification along the polarization axis. **d**, In contrast, cells treated with nocodazole, which adopt a rounded morphology, exhibited similar collagen density in all directions, both near and far from the cell, indicating isotropic matrix densification. In panel **c**, $n = 6$ for each group. The height of the bars and the error bars indicate the mean and the standard error, respectively. Statistical analysis was performed using the two-sided unpaired Student's t-test. The data were taken from our previously published work.[2]

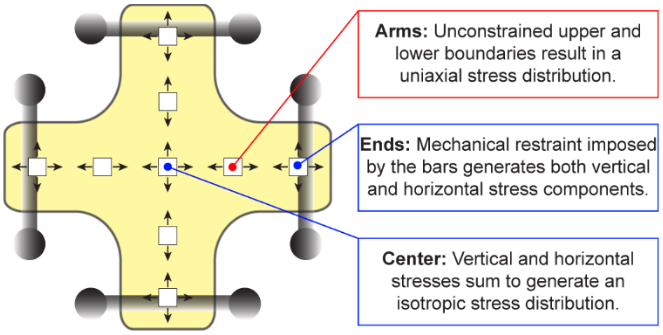


**Figure S2. Schematic representation of stress distribution in cruciform tissue constructs.** Cells contract against the anchoring bars, generating mechanical tension due to constrained contraction. In the mid-arm regions, stress is primarily aligned along the arm's longitudinal axis, producing an anisotropic stress field. At the central intersection of the specimen, equal perpendicular stress components from each arm superimpose, resulting in equal tension in both directions with no dominant stress orientation. Near the anchoring bars, vertical contraction is partially resisted by the constraint imposed by the bars, leading to reduced stress anisotropy relative to the mid-arm regions.

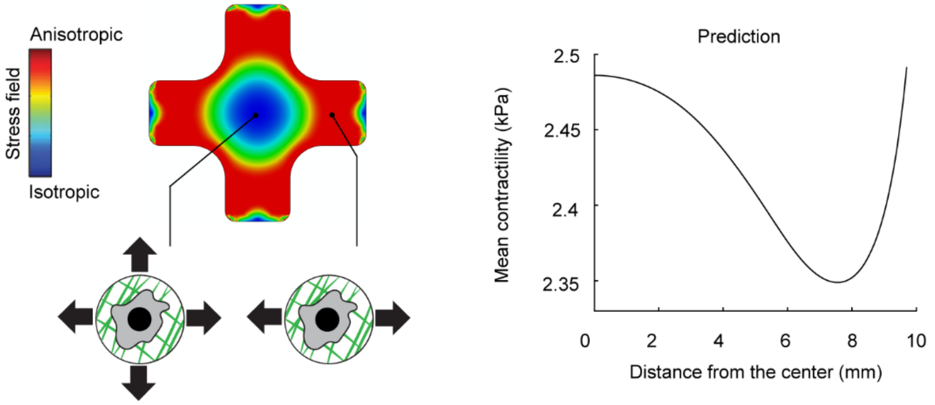


**Figure S3. Eliminating tension anisotropy from the model led to an opposite trend in predicted activation patterns compared to experimental observations.** We conducted the cruciform tissue simulations again after removing the anisotropy contribution by setting the anisotropy feedback parameter $f_a$ in Eq. (1) to zero. Under these conditions, the model predicted a spatial activation pattern opposite to the experimental results: fibroblast activation was highest in regions experiencing isotropic stress (the tissue center and near the arm ends) and lowest in regions of anisotropic tension (the mid-arm regions). Without the anisotropy term, the model indicated that fibroblasts could no longer respond to directional stress cues and instead activated solely in proportion to the mean tension magnitude. The data were taken from our previously published work.[2]

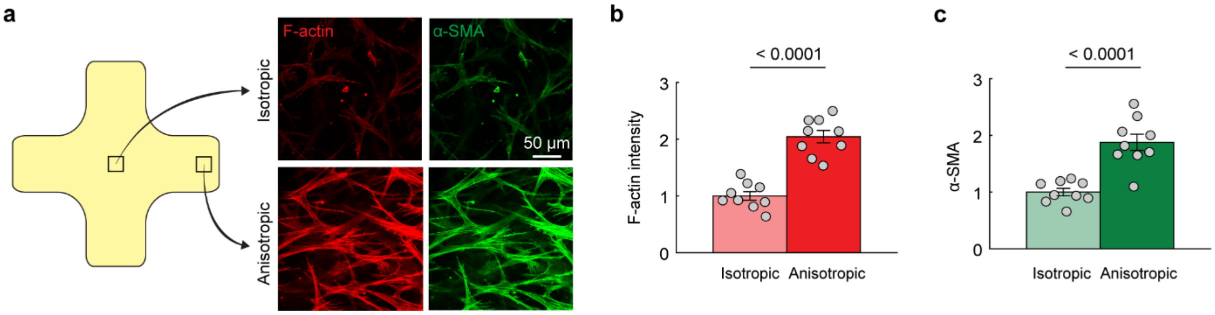


**Figure S4. Tension anisotropy enhances fibroblast activation. a**, Contraction of the cruciform-shaped tissue, anchored at the ends of its four arms, generated equibiaxial isotropic tension at the tissue center and anisotropic tension along the arms. **a-c**, Fibroblasts exposed to anisotropic stress fields in the arm regions exhibited significantly higher levels of F-actin and α-SMA, indicating increased activation toward a myofibroblast phenotype. The data and images were taken from our previously published work.[2] In panel **b-c**, $n$ = 9 for each group. The height of the bars and the error bars indicate the mean and the standard error, respectively. Statistical analysis was performed using the two-sided unpaired Student's t-test.

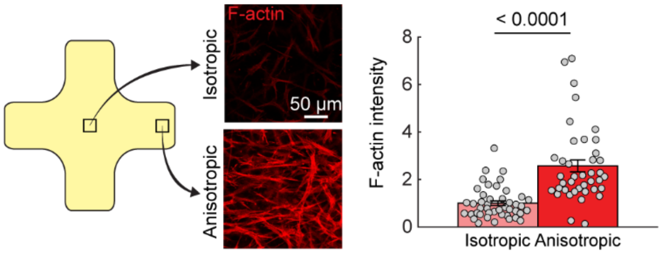


**Figure S5. Fibroblast activation promotion induced by tension anisotropy is independent of extracellular collagen concentration.** To assess whether the effect of tension anisotropy on fibroblast activation depends on collagen density in the matrix, we increased collagen concentration from 1 mg/ml (as used in Fig. 2) to 2 mg/ml. Similar to tissues with lower collagen concentration, fibroblasts embedded in higher collagen concentration exhibited significantly greater activation in anisotropic regions compared to isotropic regions, indicating that tension anisotropy enhances fibroblast activation regardless of matrix collagen concentration. The data and images were taken from our previously published work.[2] $n$ = 44 for each group. The height of the bars and the error bars indicate the mean and the standard error, respectively. Statistical analysis was performed using the two-sided unpaired Student's t-test.

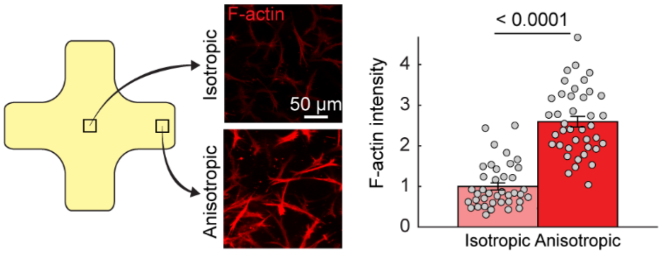


**Figure S6. Tension anisotropy enhances fibroblast activation even in the presence of matrix metalloproteinase inhibition.** To determine whether fibroblast activation induced by tension anisotropy depends on matrix degradation, fibroblasts were treated with GM6001, a well-known matrix metalloproteinase (MMP) inhibitor. Similar to untreated control cells (Fig. 2), fibroblasts exposed to anisotropic stress fields exhibited significantly higher activation levels in anisotropic regions compared to isotropic regions, despite MMP inhibition, indicating that fibroblast activation induced by tension anisotropy does not depend on matrix degradation. The data and images were taken from our previously published work.[2] $n$ = 40 for each group. The height of the bars and the error bars indicate the mean and the standard error, respectively. Statistical analysis was performed using the two-sided unpaired Student's t-test.

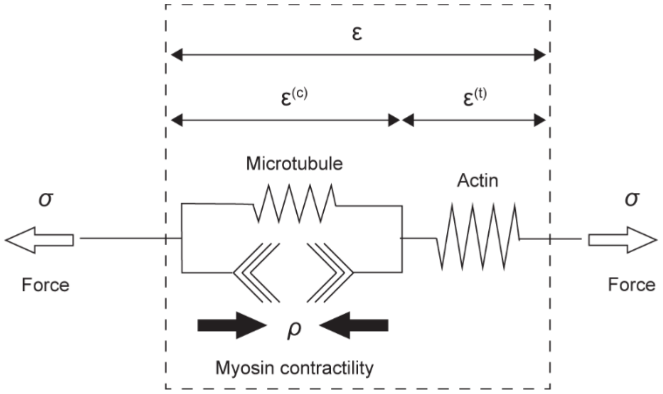


**Figure S7.** One-dimensional representation of $\rho$, $\sigma$, $\varepsilon^{(c)}$, $\varepsilon^{(t)}$, and $\varepsilon$ in our three-dimensional model.

#### Supplementary Tables

Table S1 Parameters for the theoretical cell model

| Parameter | Description | Value | Unit |
|---|---|---|---|
| $\rho_0$ | Baseline contractility | 5 | kPa |
| $\alpha_\mathrm{v}$ | Volumetric feedback parameter | 2.2 | 1/kPa |
| $\alpha_\mathrm{d}$ | Deviatoric feedback parameter | 2.2 | 1/kPa |
| $\beta_\mathrm{v}$ | Volumetric chemical stiffness parameter | 2.4 | 1/kPa |
| $\beta_\mathrm{d}$ | Deviatoric chemical stiffness parameter | 2.4 | 1/kPa |
| $E_\mathrm{MT}$ | Stiffness of microtubule | 10 | kPa |
| $\nu_\mathrm{MT}$ | Poisson's ratio of microtubule | 0.4 | |
| $E_\mathrm{A}$ | Stiffness of actin | 10 | kPa |
| $\nu_\mathrm{A}$ | Poisson's ratio of actin | 0.4 | |
| $\alpha_\mathrm{a}$ | Tension anisotropy parameter | 1.0 | |
| $l$ | Actin network strain-stiffening parameter | 50 | |
| $m$ | Actin network strain-stiffening parameter | 4.0 | |
| $n$ | Actin network strain-stiffening parameter | 4.0 | |
| $\epsilon_\mathrm{c}$ | Critical tensile principal strain of actin network | 0.1 | |
